# *Mcidas* mutant mice reveal a two-step process for the specification and differentiation of multiciliated cells in mammals

**DOI:** 10.1101/439703

**Authors:** Hao Lu, Priyanka Anujan, Feng Zhou, Yiliu Zhang, Yan Ling Chong, Colin D. Bingle, Sudipto Roy

**Affiliations:** Institute of Molecular and Cell Biology, Proteos, 61 Biopolis Drive, Singapore 138673, Singapore; Academic Unit of Respiratory Medicine, Department of Infection, Immunity and Cardiovascular Disease, University of Sheffield, Sheffield S10 2JF, UK; Department of Pediatrics, Yong Loo Lin School of Medicine, National University of Singapore, 1E Kent Ridge Road, Singapore 119288; Department of Biological Sciences, National University of Singapore, 14 Science Drive 4, Singapore 117543, Singapore

**Keywords:** cilia, multiciliated cell, GMNC, MCIDAS, E2F, deuterosome

## Abstract

Motile cilia on multiciliated cells (MCCs) function in fluid clearance over epithelia. Studies with *Xenopus* embryos and patients with the congenital respiratory disorder reduced generation of multiple motile cilia, have implicated the nuclear protein MCIDAS (MCI), in the transcriptional regulation of MCC specification and differentiation. Recently, a paralogous protein, GMNC, was also shown to be required for MCC formation. Surprisingly, and in contrast to the presently held view, we find that *Mci* mutant mice can specify MCC precursors. However, these precursors cannot produce multiple basal bodies, and mature into single ciliated cells. We show that MCI is required specifically to induce deuterosome pathway components for the production of multiple basal bodies. Moreover, GMNC and MCI associate differentially with the cell-cycle regulators E2F4 and E2F5, which enables them to activate distinct sets of target genes (ciliary transcription factor genes versus genes for basal body generation). Our data establish a previously unrecognized two-step model for MCC development: GMNC functions in the initial step for MCC precursor specification. GMNC induces *Mci* expression, which then drives the second step of basal body production for multiciliation.

**SUMMARY STATEMENT:** We show how two GEMININ family proteins function in mammalian multiciliated cell development: GMNC regulates precursor specification and MCIDAS induces multiple basal body formation for multiciliation.

## INTRODUCTION

Health of our airways is critically dependent on mucociliary clearance, a process by which pathogen- and pollutant-laden mucus is cleared out by the beating of hundreds of motile cilia that decorate the surfaces of MCCs (Bustamante-Marin and Ostrowski, 2017). Ineffective mucus clearance predisposes individuals to respiratory diseases, best exemplified by congenital disorders like primary ciliary dyskinesia (PCD) and reduced generation of multiple motile cilia (RGMC) (Knowles et al., 2016). In PCD, MCCs form normally, but their cilia are immotile or have defective motility due to mutations in proteins of the motility apparatus. By contrast, in RGMC, differentiation of multiple cilia or the MCCs themselves is affected. MCCs are also present within brain ventricles, where they drive circulation of cerebrospinal fluid as well as within reproductive organs, where they promote mixing of reproductive fluids and germ cell transportation (Zhou and Roy, 2015, Brooks and Wallingford, 2014).

Post-mitotic MCC precursors support an explosive production of numerous basal bodies that migrate to the apical surface and nucleate the biogenesis of multiple motile cilia. One key aspect of MCC development is the transcriptional program required to institute its fate and its unique differentiation program, which has just begun to be elucidated (Spassky and Meunier, 2017). Studies with *Xenopus* embryos, which differentiate epidermal MCCs for mucus clearance, have implicated a small coiled-coil Geminin family protein, Mcidas (Mci; aka Multicilin), as a key regulator of MCC fate (Stubbs et al., 2012). Morpholino-mediated inhibition of Mci function in the frog resulted in a complete loss of the MCCs, indicating an essential role of the protein for their specification and differentiation. This phenotype is largely conserved in RGMC patients carrying mutations in *MCIDAS*, encoding human MCI, with their airways populated by cells differentiating only one or two immotile cilia (Boon et al., 2014). Current evidence suggests that on the one hand MCI regulates transcription of genes encoding transcription factors (such as FOXJ1) that activate genes for ciliary differentiation and motility, and on the other genes for the production of multiple basal bodies (such as *Ccno*, *Deup1*, *Cep152* and *Ccdc78*) (Ma et al., 2014). MCI lacks a DNA binding domain, and is thought to regulate transcription by associating with the cell-cycle regulators E2F4 or E2F5, and their obligatory dimerization partner DP1 (Ma et al., 2014).

Recently, another MCI-related protein, GMNC (aka GEMC1), has been identified as an essential regulator of MCC development (Zhou et al., 2015, Terré et al., 2016, Arbi et al., 2016). Zebrafish and mice with mutations in *Gmnc* completely lack MCCs. Although there is disagreement on whether GMNC functions with the E2F proteins (Zhou et al., 2015, Terré et al., 2016), nevertheless, like MCI, it can fully activate the transcriptional program for MCC specification and differentiation. Both *Mci* and *Gmnc* are expressed quite specifically in developing MCCs: GMNC acts upstream of MCI and it is required for *Mci* expression in MCC precursors, whereas MCI is unable to induce *Gmnc* (Arbi et al., 2016, Terré et al., 2016, Zhou et al., 2015). What remains presently unclear is how two related proteins, with purported similar transcriptional activities, can have near identical effects on the MCC developmental program. Given this quandary, we re-investigated *Mci* function, this time by stably inactivating the gene in mice. We now show that in contrast to the presently held belief that MCI regulates MCC specification as well as differentiation, *Mci* mutant mice can specify MCC precursors in normal numbers, which express a suite of genes for the transcriptional regulation of ciliary differentiation. However, these cells are unable to activate genes for basal body production, and consequently, differentiate single motile-like cilia. Moreover, we show that while MCI interacts with E2F4 and E2F5, GMNC forms a complex preferentially with E2F5, with distinct C-terminal domains of the two proteins determining this differential interaction. We argue that MCC precursor specification and induction of transcription factors for ciliary gene expression is regulated by GMNC. In the next step, MCI amplifies the expression of ciliary transcription factors and triggers the expression of genes required for biogenesis of multiple basal bodies. These basal bodies then seed the assembly of multiple cilia to complete the process of MCC differentiation. Thus, our study provides mechanistic insight into how the regulatory activities of two paralogous proteins coordinately organize the transcriptional program of a specialized ciliated cell-type.

## RESULTS

### *Mci* mutant mice cannot differentiate MCCs with multiple cilia

We used the CRISPR/Cas9 technology to generate a mutant allele of mouse *Mci*. This allele, a deletion of 32 bp in exon 2 of the *Mci* gene, is predicted to encode a severely C-terminally truncated MCI protein, lacking all of the important functional domains (coiled-coil domain in the middle of the protein and the TIRT domain for E2F/DP1 interaction at the C-terminus (Ma et al., 2014)), implying a strong loss-of-function condition (see methods and Fig. S1A-D). Heterozygous mice exhibited no phenotypic abnormalities, and when intercrossed, homozygous wild-type, heterozygous as well as homozygous mutants were recovered in the correct Mendelian ratio. However, the homozygous mutants were runted compared to their wild-type and heterozygous siblings, and showed progressive post-natal lethality (Fig. S2A-C). Since *Gmnc* mutant mice also exhibit similar phenotypes, and their lethality was attributed to the development of hydrocephalus (Terré et al., 2016), we examined the *Mci* mutants for this defect. Indeed, histological analysis of the brains of two mutant animals (n = 2) showed hydrocephalus, suggestive of dysfunctional ependymal MCCs (data not shown, but see Fig. S2D,E). Moreover, all homozygous mutants tested (males and females) failed to breed, when in-crossed as well as when out-crossed, indicating MCC defects in the reproductive organs. To adduce evidence that the production of the wild-type MCI protein is indeed disrupted in the homozygotes, we cloned the mutant *Mci* cDNA from tracheal tissue and confirmed the presence of the 32 bp deletion (Fig. S1E). Moreover, quantitation of *Mci* transcript levels from cultures of airway cells from the homozygotes revealed severe reduction relative to wild-type (see Fig. 4 below).

We next investigated the status of MCCs in tissues where they are normally known to differentiate – trachea, oviducts and brain ependyma (Brooks and Wallingford, 2014, Spassky and Meunier, 2017, Zhou and Roy, 2015). In the wild-type, abundant MCCs with multiple motile cilia were visible decorating the luminal surface of these tissues, interspersed with other cell-types (Fig. 1A,B and data not shown). By contrast, in the mutants we found a complete loss of the multiple ciliated cells (Fig. 1C,D and data not shown). Instead, we could observe cells with a single cilium. The length and width of these monocilia were similar to the multiple cilia of MCCs, but distinctly different from the shorter, thinner primary cilia present on neighboring cells (Fig. 1C). Even though the *Mci* mutants develop hydrocephalus and are infertile, because these mice are maintained under specific-pathogen-free (SPF) conditions, we did not detect any obvious symptoms of airway disease either at the behavioral level or through histological analysis of respiratory tissues (data not shown).

**Fig. 1.**
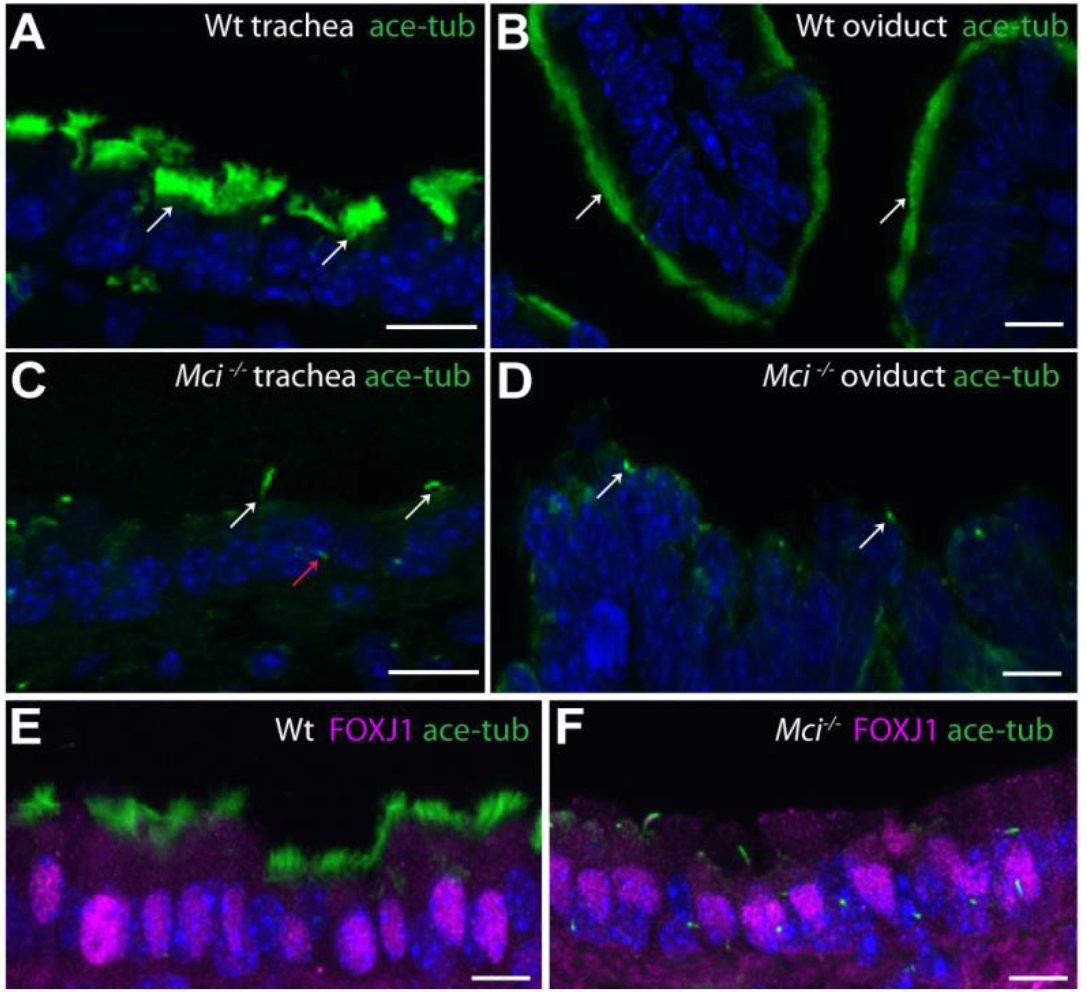
MCCs in *Mci* mutant mice differentiate a single cilium and express FOXJ1. (A) Wild-type trachea section showing multiple cilia on MCCs (arrows). (B) Wild-type oviduct section showing multiple cilia on MCCs (arrows). (C) *Mci* mutant trachea section showing cells with single cilium (white arrows). A primary cilium in a neighboring cell is indicated (red arrow). (D) *Mci* mutant oviduct section showing cells with single cilium (arrows). (E) Nuclear localized FOXJ1 expression in MCCs of wild-type trachea. (F) Nuclear localized FOXJ1 expression in monociliated cells of *Mci* mutant trachea. In all preparations, cilia were stained with anti-acetylated tubulin antibodies (green) and nuclei with DAPI (blue). Wt, wild-type. Scale bars, 10 μm. For all histological data presented in this and other figures, tissues from at least 2 wild-type and 3 *Mci* mutant mice were analyzed, unless otherwise mentioned.

### In *Mci* mutants, MCC precursors are specified but fail to generate multiple basal bodies

To begin to uncover the developmental defect underlying MCC absence in *Mci* mutants, we first analyzed the expression of FOXJ1, a protein that is required to activate the motile cilia-specific transcriptional program (Yu et al., 2008, Stubbs et al., 2008, Choksi et al., 2014a). Previous studies with *Xenopus* embryos and human airway cells have shown that *Foxj1* is a transcriptional target of MCI (Boon et al., 2014, Stubbs et al., 2012). Strikingly, and contrary to these earlier findings, FOXJ1 expression was not affected in MCC harboring tissues of *Mci* mutant mice. While FOXJ1 was present in the nucleus of wild-type MCCs, in the mutants, we found nuclear-localized FOXJ1 in cells bearing single long cilium (Fig. 1E,F and Fig. S2D,E). Based on this observation, we reasoned that the absence of MCI function perhaps does not compromise the specification of MCC precursors, but instead is required in these cells to differentiate multiple cilia. To bolster this view, we analyzed the expression of a suite of additional transcription factors, which, like FOXJ1, have been implicated in motile ciliogenesis: RFX2, RFX3 and TAP73 (Choksi et al., 2014b, Jackson and Attardi, 2016). Again, like FOXJ1, expression of these transcription factors was not appreciably affected (Fig. 2A-F). These findings suggest that in the absence of MCI function, MCC precursors get specified normally, but they then differentiate a single cilium instead of multiple cilia. To garner evidence that this single cilium possesses attributes of motile cilia, we stained tracheal sections with antibodies against RSPH1 and RSPH9, two radial spoke-head proteins that are unique structural components of motile cilia (Frommer et al., 2015). The single cilium of *Mci* mutants showed localization of both these proteins along the axoneme (Fig. 2G-J and Fig. S3A-D). We also examined the status of the MCCs in the trachea using scanning electron microscopy (SEM). Unlike in the wild-type, where hundreds of motile cilia were present at the apical surface of the MCCs, *Mci* mutant trachea showed cells with a single cilium whose dimensions were similar to the individual motile cilium of the wild-type MCCs (Fig. 2K,L).

**Fig. 2.**
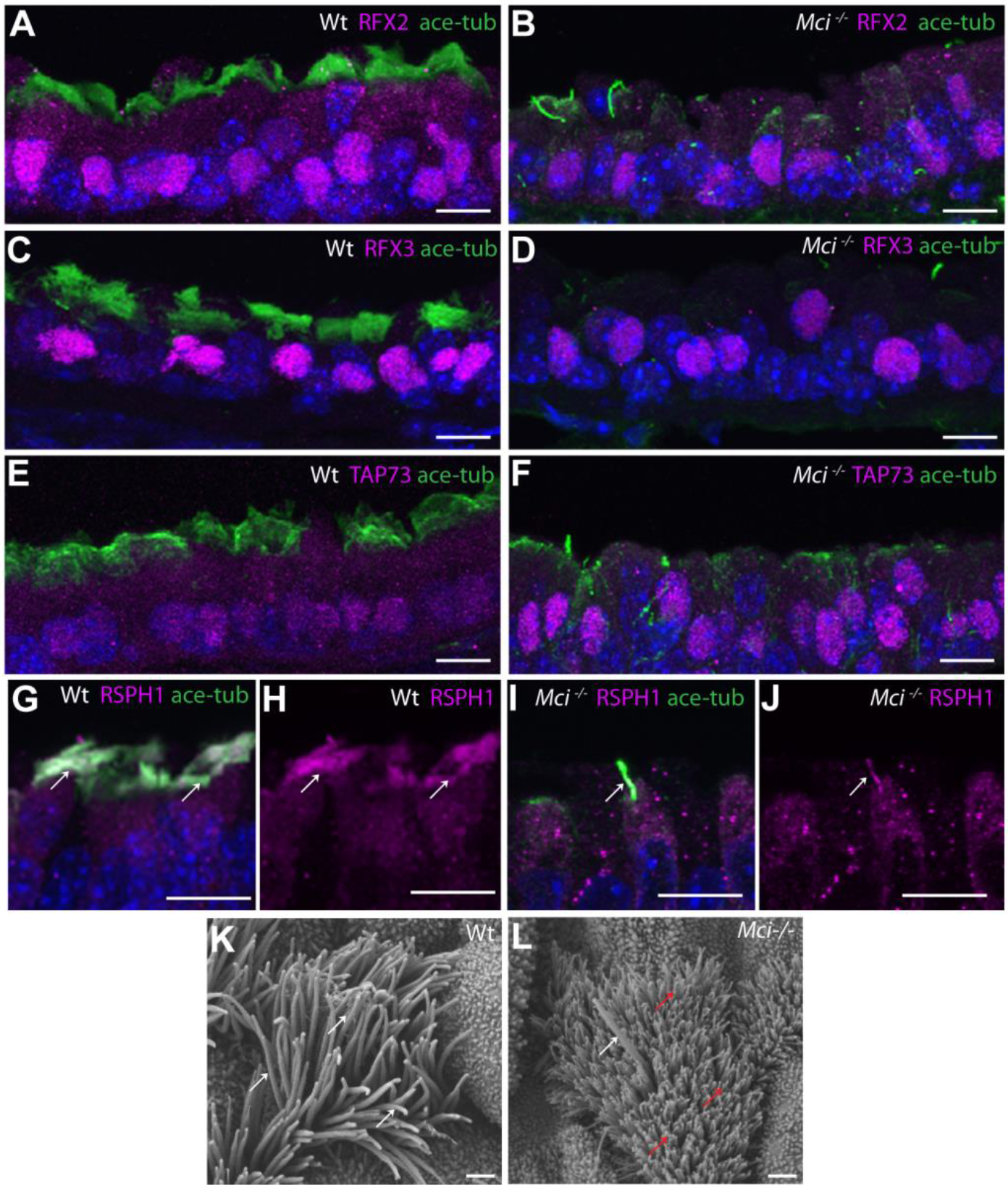
*Mci* mutant MCCs precursors express a suite of ciliary transcription factors and their single cilium localizes motile cilia-specific proteins. (A) Nuclear localized RFX2 expression in MCCs of wild-type trachea. (B) Nuclear localized RFX2 expression in monociliated cells of *Mci* mutant trachea. (C) Nuclear localized RFX3 expression in MCCs of wild-type trachea. (D) Nuclear localized RFX3 expression in monocilated cells of *Mci* mutant trachea. (E) Nuclear localized TAP73 expression in MCCs of wild-type trachea. (F) Nuclear localized TAP73 expression in monociliated cells of *Mci* mutant trachea. (G) RSPH1 co-localization with acetylated tubulin to MCC cilia of wild-type trachea (arrows). (H) RSPH1 localization to MCC cilia of wild-type trachea (arrows; display of only RPSH1 staining from panel G). (I) RSPH1 co-localization with acetylated tubulin to single cilium of *Mci* mutant trachea (arrow). (J) RSPH1 localization to single cilium of *Mci* mutant trachea (arrow; display of only RSPH1 staining from panel I). (K) SEM analysis of a wild-type tracheal MCC showing multiple cilia (arrows). (L) SEM analysis of *Mci* mutant MCC with a single cilium (white arrow). The microvilli, which are quite long in the MCCs and normally remain obscured by the multiple cilia, are indicated (red arrows). One wild-type and one mutant trachea were scanned by SEM. The single cilium phenotype of the *Mci* mutant trachea is representative of several fields of view scanned by SEM. In all preparations, cilia were stained with anti-acetylated tubulin antibodies (green) and nuclei with DAPI (blue). Scale bars, A-J = 10 μm; K,L = 5 μm.

Since MCC differentiation is contingent upon the generation of multiple basal bodies, we next investigated the status of these organelles. Staining with anti-PERICENTRIN antibodies revealed multiple basal bodies in wild-type MCCs, neatly arrayed along their apical membranes (Fig. 3A). By contrast, in *Mci* mutants, we could observe a single basal body associated with the single cilium (Fig. 3B). We obtained similar data with antibodies to γ-tubulin, which also labels ciliary basal bodies (data not shown; but see next section). Thus, the program for multiple basal body generation is significantly derailed in *Mci* mutants. We used transmission electron microscopy (TEM) to analyze the basal body phenotype in greater subcellular detail. While cilia-bearing multiple basal bodies were readily visible in wild-type MCCs, in the mutants, they were absent from several sections that we examined (Fig. 3C,D).

**Fig. 3.**
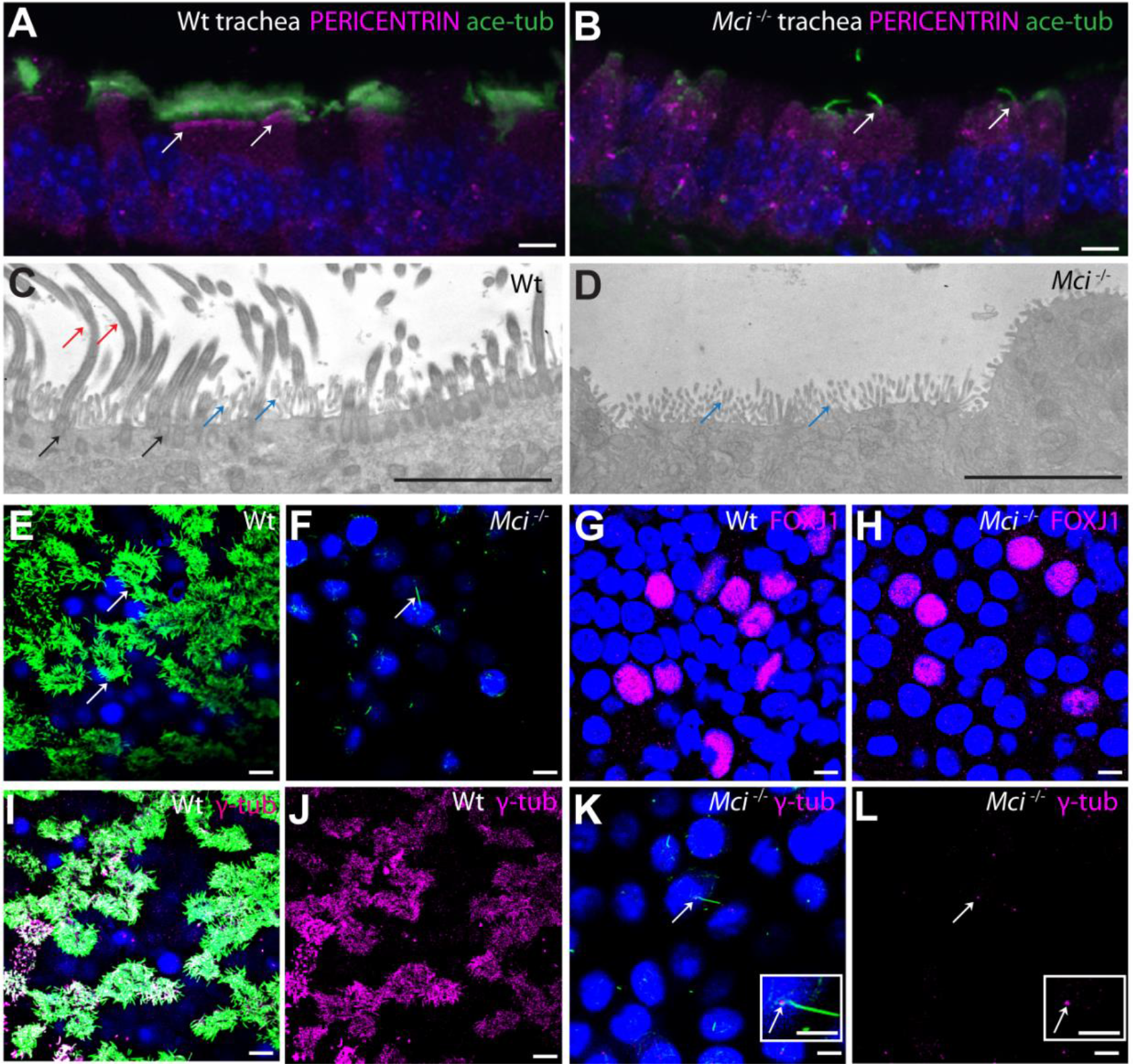
*Mci* mutant MCC precursors are unable to generate multiple basal bodies. (A) Wild-type trachea section showing apically aligned multiple basal bodies in MCCs (arrows; stained with anti-PERICENTRIN antibodies). (B) Section of *Mci* mutant trachea showing single basal body in monociliated cells (arrows). (C) TEM image showing multiple basal bodies (black arrows) and cilia (red arrows) in a wild-type MCC. Microvilli are also indicated (blue arrows). (D) TEM image showing lack of multiple basal bodies and cilia in a *Mci* mutant MCC. Microvilli are indicated (blue arrows). 5 sections each from 2 independent wild-type and mutant tracheae were sampled. (E) Wild-type MCCs differentiated in ALI culture with multiple cilia (arrows). (F) *Mci* mutant airway cells differentiated in ALI culture with single cilium (arrow). (G) Wild-type airway cells differentiated in ALI culture showing nuclear FOXJ1 expression. (H) *Mci* mutant airway cells differentiated in ALI culture showing nuclear FOXJ1 expression. (I) Wild-type MCCs differentiated in ALI culture with multiple basal bodies (stained with anti-γ-tubulin antibodies) and multiple cilia. (J) Display of only γ-tubulin staining from panel I. (K) *Mci* mutant cells differentiated in ALI culture with single basal body (arrow) and single cilium. Inset shows single cilium and basal body (arrow). (L) Display of only γ-tubulin staining from panel K showing single basal body (arrow) Inset shows single basal body (arrow). In preparations shown in panels A,B,E,F,I,K cilia were stained with anti-acetylated tubulin antibodies (green) and nuclei were stained with DAPI (blue). Scale bars, 5μm. ALI cultures were done in 3 independent biological replicates.

### *In vitro* culture of *Mci* mutant airway cells revealed a strong impairment in expression of basal body generation genes

To uncover the earliest developmental defects in *Mci* mutant MCCs, we resorted to culture of mouse tracheal epithelial cells (mTECs) *in vitro*, followed by differentiation under air-liquid interface (ALI) condition. Consistent with our observations from tracheal sections, mTECs from *Mci* mutant mice differentiated single cilium bearing cells, unlike the wild-type where MCCs readily formed (Fig. 3E,F). Moreover, expression of FOXJ1 was not affected in *Mci* mutant cultures (Fig. 3G,H), implying that like *in vivo*, loss of MCI does not compromise the ability to adopt the MCC precursor identity (RSPH proteins also localized to the single cilium of ALI cultured cells (data not shown)). Moreover, γ-tubulin staining revealed absence of multiple basal bodies, whereas wild-type MCCs showed clouds of basal bodies at their apical surface (Fig. 3I-L). We obtained a similar result with antibodies against the CENTRIN protein that also marks the basal bodies (Fig. S3E-H). Thus, *in vivo* as well as *in vitro*, loss of MCI specifically compromises the ability of MCC precursors to generate multiple basal bodies.

We next used RT-qPCR analysis to interrogate the transcriptional profile of the *Mci* mutant cells. While *Mci* transcripts were strongly reduced, expression of genes encoding upstream transcription regulatory factors such as GMNC, FOXJ1 and the RFX family members RFX2 and RFX3, were not appreciably affected or were slightly reduced relative to wild-type (Fig. 4A-E). This lack of a major reduction is consistent with our observations using immunofluorescence analysis, described above. By contrast, genes implicated in the production of multiple basal bodies – *Deup1*, *Ccdc78*, *Ccno* and *Cdc20b* - were all strongly reduced, indicating that MCI is specifically required to activate their transcription (Fig. 4F-I).

**Fig. 4.**
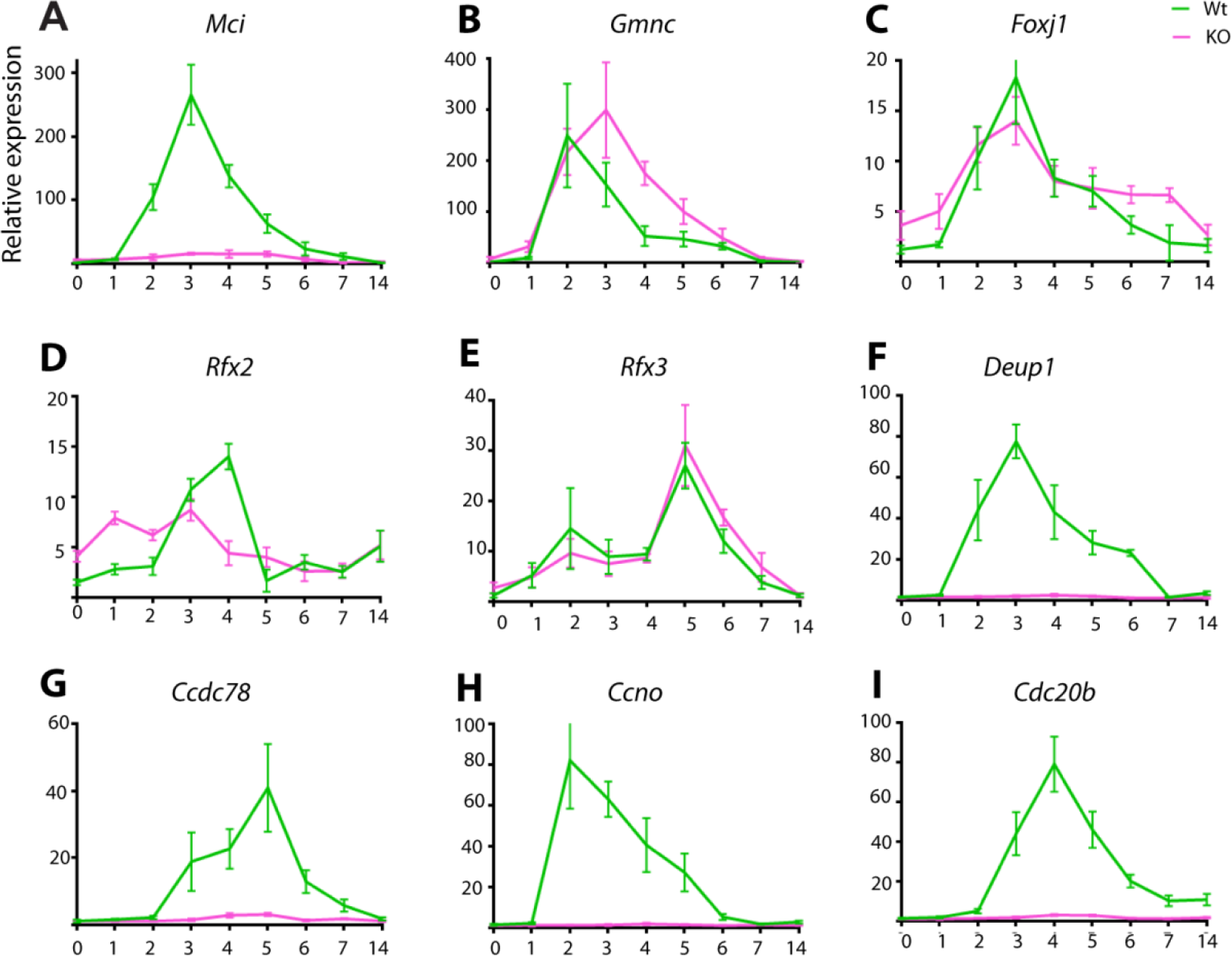
RT-qPCR analysis of ciliary transcription factor and DD pathway genes expression levels between wild-type and *Mci* mutant airway cells in ALI culture. (A-I) Relative expression levels have been plotted along the *y*-axis, and days in ALI culture along the *x*-axis. KO = *Mci* mutant. Error bars: standard error of the mean (SEM). Analysis was done on 3 independent biological replicates.

### In *Mci* mutant MCC precursors, deuterosomes are severely reduced in numbers and are dysfunctional

Current view posits two distinct pathways for multiple basal body generation in MCCs. Some of the basal bodies are believed to be generated by the mother centriole-dependent (MCD) pathway, through the activity of proteins like CEP63, CEP152, PLK4 and SAS6, which also function in centriole duplication during regular cell division (Al Jord et al., 2014, Spassky and Meunier, 2017). In addition to this, a dedicated pathway exists in the MCCs for basal body generation. The vast majority of basal bodies are produced by this alternative mechanism – the deuterosome-dependent (DD) pathway – in which DEUP1, CCNO, CCDC78 and CDC20B are believed to be dedicated components (Spassky and Meunier, 2017, Klos Dehring et al., 2013, Zhao et al., 2013, Funk et al., 2015, Revinski et al., 2017). Here, electron dense structures called deuterosomes are first generated by the oligomerization of the CEP63 paralog, DEUP1. Although, whether the deuterosomes are nucleated by existing centrioles or arise *de novo* is presently a matter of debate (Al Jord et al., 2014, Zhao et al., 2018), what is clear is that after formation, they recruit CEP152 and other MCD pathway proteins (PLK4, SAS6 etc) to generate multiple procentrioles. These procentrioles then mature into centrioles, detach and migrate apically to dock with the plasma membrane and form ciliary basal bodies.

We found that despite the strong reduction in *Deup1* mRNA levels in *Mci* mutants, DEUP1-positive deuterosomes nevertheless formed, albeit in severely reduced numbers (Fig. 5A-E). However, since we consistently failed to detect multiple centrioles in *Mci* mutant MCCs *in vitro* as well as *in vivo*, these must be defective deuterosomes incapable of supporting centriole biogenesis. The complete absence of centriole duplication in *Mci* mutants suggests that even the MCD pathway is defective. To investigate this issue further, we examined expression of the MCD pathway gene *Cep63*, as well as *Cep152*, *Plk4* and *Sas6* (which are shared by both DD and MCD pathways), but failed to detect major differences in their expression levels (Fig. 5F-I).

**Fig. 5.**
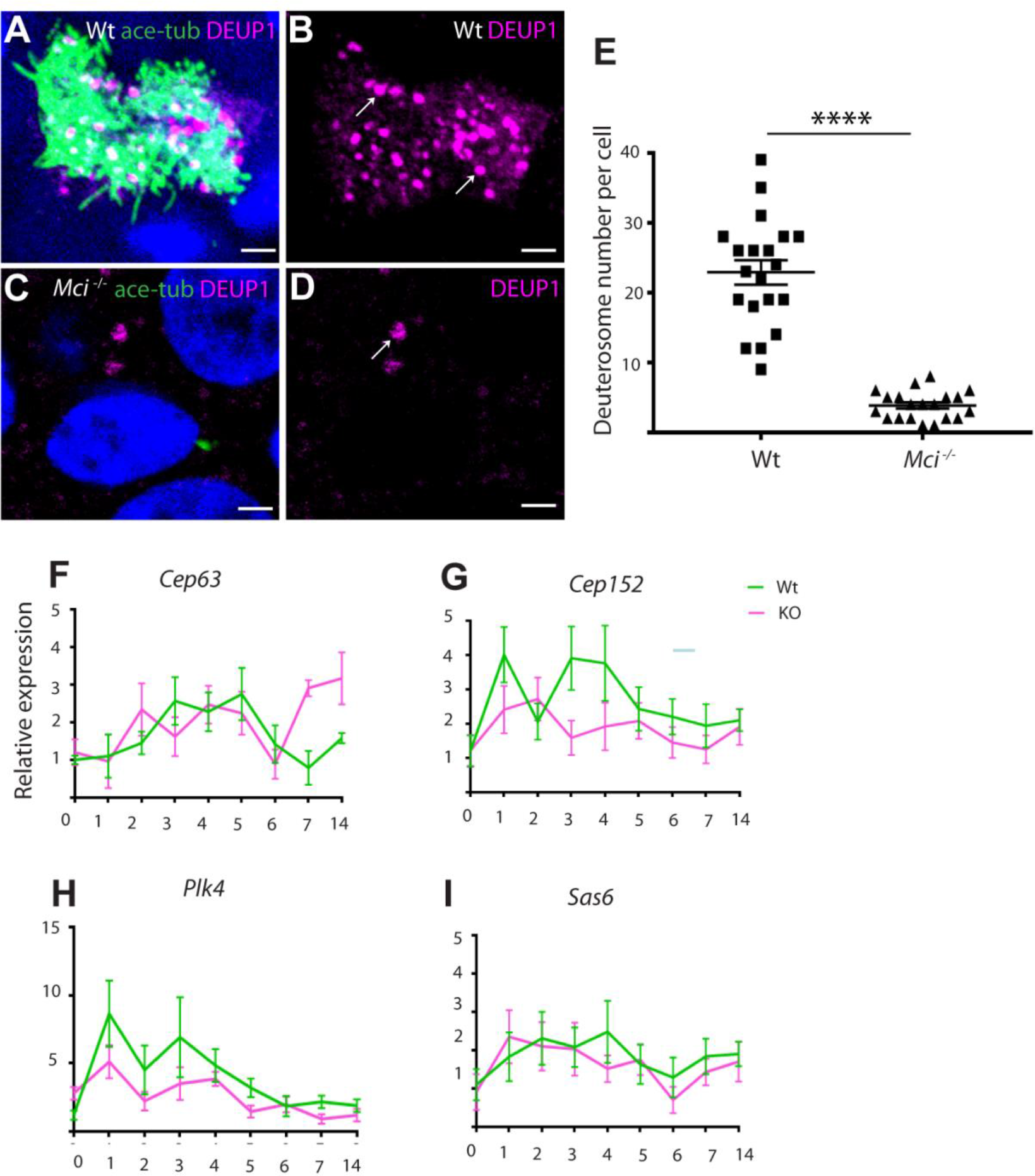
In *Mci* mutants, the DD pathway for basal body production is strongly affected but not the MCD pathway. (A) ALI cultured wild-type MCCs showing DEUP1-positive deuterosomes. (B) Display of only DEUP1 staining from panel A, showing deuterosomes (arrows). (C) ALI cultured *Mci* mutant airway cells showing DEUP1-positive deuterosomes. (D) Display of only DEUP1 staining from panel C, showing a deuterosome (arrow). Scale bars, 5 μm. (E) Quantification of numbers of DEUP1+ deuterosomes in differentiating wild-type and *Mci* mutant MCCs under ALI conditions. 20 cells were counted for each genotype at ALI day 3. p: **** ≤ 0.0001. (F-I) RT-qPCR analysis of MCD pathway gene expression levels between wild-type and *Mci* mutant airway cells in ALI culture. Relative expression levels have been plotted along the *y*-axis, and days in ALI culture along the *x*-axis. Error bars: SEM. Analysis was done on 3 independent biological replicates.

### GMNC and MCI have distinct effects on the MCC-specific transcriptional program

In *Gmnc* mutant mice, the expression of the entire MCC-specific transcriptional program is significantly dampened (Terré et al., 2016). This includes (i) genes for ciliary transcription factors like FOXJ1 and MCI as well as (ii) genes for DD (but not MCD) pathway proteins. On the other hand, our current analysis shows that MCI loss preferentially affects the DD pathway genes. To examine this differential effect, we first over-expressed the human homologs of GMNC and MCI individually in HEK293T cells, and monitored the expression of genes from the two sets mentioned above. Terré et al. have previously demonstrated that HEK293T cells can be used effectively to assess the transcriptional activities of GMNC and MCI (Terré et al., 2016). GMNC could induce *MCI*; however, over-expression of MCI could not induce *GMNC* (Fig. 6A,B), which is consistent with previous reports (Arbi et al., 2016, Terré et al., 2016). Interestingly, both GMNC and MCI were able to induce *FOXJ1* (Fig. 6C). With respect to DD pathway genes, MCI alone or MCI together with GMNC strongly up regulated *DEUP1*, *CCNO* and *CDC20B* (Fig. 6D-F), although there was no obvious additive effect from the co-expression. Whereas, GMNC alone could only weakly induce these genes (Fig. 6D-F). These data support the idea that MCI preferentially affects the expression of DD pathway genes (also see below).

**Fig. 6.**
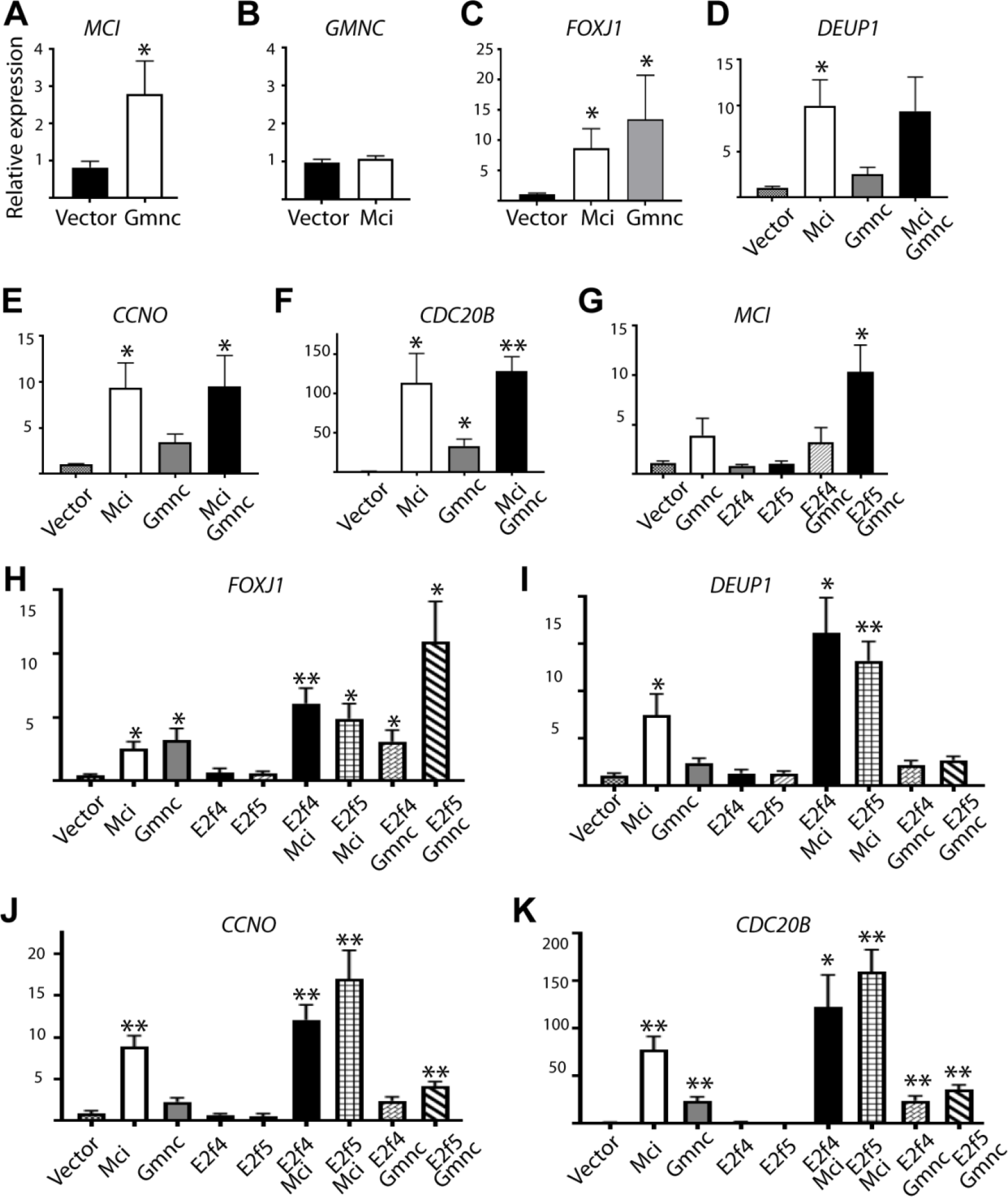
RT-qPCR analysis of ciliary transcription factor and DD pathway genes expression levels on over-expression of MCI, GMNC and E2F proteins in HEK293T cells. (A-K) Relative expression levels have been plotted along the *y*-axis, and over-expression conditions indicated along the *x*-axis. Error bars: SEM. Analysis was done on 3 independent biological replicates. p: * ≤ 0.05, ** ≤ 0.01.

Since MCI, and also GMNC, have been reported to interact with E2F4 and E2F5 for transcription, we next checked the transcriptional abilities of GMNC and MCI when co-expressed with E2F4 or E2F5. The ability of GMNC to induce *MCI* and *FOXJ1* was strongly increased with E2F5, but not with E2F4 (Fig. 6G,H). Likewise, a slight upregulation of DD pathway genes occurred when GMNC was over-expressed with E2F5, but not with E2F4 (Fig. 6I-K). By contrast, MCI with E2F4 or E2F5 had stronger transcriptional effect on *DEUP1, CCNO* and *CDC20B* than MCI alone (Fig. 6I-K). With regard to *FOXJ1*, MCI with E2F4 as well as E2F5 could induce higher levels of transcription than MCI alone, and MCI with E2F4 was more efficient than MCI with E2F5 (Fig. 6H). Thus, the transcriptional activity of GMNC appears to be much more effective with E2F5, whereas MCI regulates its target genes with either E2F4 or E2f5.

### Differential interaction of E2F4 and E2F5 with MCI and GMNC

We previously showed that human GMNC is unable to interact effectively with E2F4 (Zhou et al., 2015). However, Terré et al. demonstrated that GMNC can interact with E2F4 as well as E2F5 (Terré et al., 2016). Moreover, E2F5 has been previously shown to significantly potentiate the transcriptional activity of GMNC (Arbi et al., 2016). Since our current analysis shows that GMNC and MCI act in a step-wise manner and regulate distinct sets of targets, we reevaluated their interactions with the E2F factors. Consistent with our earlier report, human as well as mouse GMNC interacted poorly, if at all, with human and mouse E2F4 and DP1, respectively (Fig. 7A,B). By contrast, we found robust interaction of human and mouse GMNC with human and mouse E2F5 and DP1, respectively (Fig. 7A,B). On the other hand, as reported before (Ma et al., 2014), MCI proteins from both species interacted equally efficiently with E2F4 and E2F5 (Fig. 7A and Fig. S4A).

**Fig. 7.**
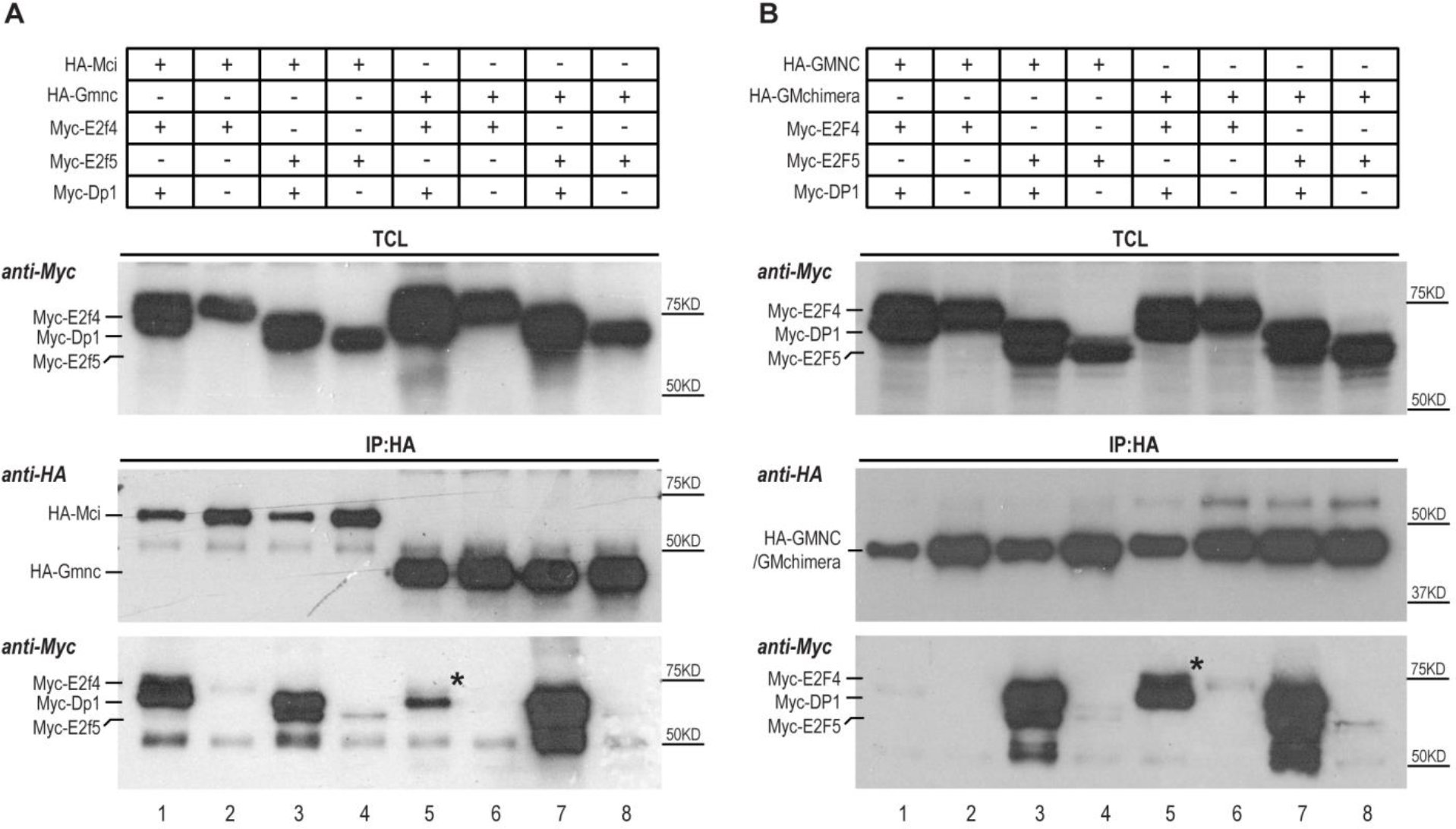
Differential interaction of MCI and GMNC with E2F4 and E2F5. (A) While MCI effectively interacted with both E2F4 and E2F5 in the presence of DP1 (lane 1, IP panel, anti-Myc blot), very little E2F4 co-precipitated with GMNC and DP1 (lane 5, IP panel, anti-Myc blot). The weak presence of E2F4 in the GMNC IP is marked by the black asterisk. Mouse proteins were used for this experiment. (B) C-terminal domain of MCI accounts for effective interaction with both E2F4 and E2F5. GMNC showed minimal interaction with E2F4 (lane 1, IP panel, anti-Myc blot). In comparison, the GM (engineered by replacing the C-terminus domain of GMNC with that of MCI) chimera could co-precipitate with both E2F4 and E2F5, in the presence of DP1 (lane 5 and 7, IP panel, anti-Myc blot). The black asterisk marks the E2F4 band that is absent in lane 1 (IP panel, anti-Myc blot). Human proteins were used for this experiment. The E2F and DP1 proteins were tagged N-terminally with the Myc epitope and the GMNC and MCI proteins were tagged N-terminally with the HA epitope. TCL: Total cell lysate. IP: Immunoprecipitation. Data are representative of 2 independent biological replicates.

The E2F/DP1 interaction domain in GMNC and MCI is located at the C-terminal end (approximately 40 amino acids – the TIRT domain) (Ma et al., 2014, Terré et al., 2016). We replaced this domain in GMNC with the one from MCI (Fig. S4B), and then examined the interaction of the chimera (GM) with E2F4 and E2F5. Similar to MCI, but unlike wild-type GMNC, the GM chimera efficiently bound E2F4 as well as E2F5 (Fig. 7B). Despite this, GM over-expression alone or in combination with the E2F factors failed to elicit a transcriptional response in HEK293T cells, indicating that association with E2F4 by itself is not sufficient to switch the transcriptional activity pattern of GMNC towards that of MCI (Fig. S4C,D).

### MCI can substitute for GMNC, but GMNC cannot substitute for MCI in MCC formation

Lastly, we investigated whether GMNC and MCI can substitute for each other in MCC development. Since the function of GMNC in MCC formation is quite conserved between zebrafish and mice (Arbi et al., 2016, Terré et al., 2016, Zhou et al., 2015), we first over-expressed mouse MCI in *gmnc* mutant zebrafish embryos, which completely lack MCCs from all MCC bearing tissues, and found very efficient rescue of MCCs within the pronephric (kidney) ducts, where these cells promote urine flow (Fig. 8A-C). For the converse experiment, we used lentivirus-mediated human GMNC over-expression in mTEC ALI cultures from *Mci* mutant and wild-type mice. While GMNC produced ectopic MCCs in the wild-type, it failed to rescue MCC development in *Mci* mutant cultures (Fig. 8D-G and Fig. S5A,B,D). Moreover, while GMNC over-expression in *Mci* mutant cells could induce *Mci*, *Foxj1* and *Rfx3*, DD pathway genes were not upregulated at all (Fig. S6A,B,D-F and data not shown). This observation suggests that the weak induction of DD pathway genes on over-expression of GMNC in HEK293T cells that we noted earlier (cf. Fig. 6D-F), must occur via GMNC-dependent induction of MCI. By contrast, human MCI over-expression generated significant numbers of MCCs in wild-type as well as *Mci* mutant cultures, denoting effective rescue, and also induced high levels of *Foxj1* and DD pathway genes (Fig. 8H,I and Figs. S5A,C,D and S6C-F). Both the human *GMNC* and *MCI* genes were clearly over-expressed in these experiments (Fig. S5E,F), so the inability of GMNC to rescue is unlikely to be due to inadequate over-expression. Moreover, *E2f4*, *E2f5* and *Dp1* levels were also not affected in *Mci* mutant cells, and therefore, cannot also account for the lack of rescue of MCC formation by GMNC (data not shown).

**Fig. 8.**
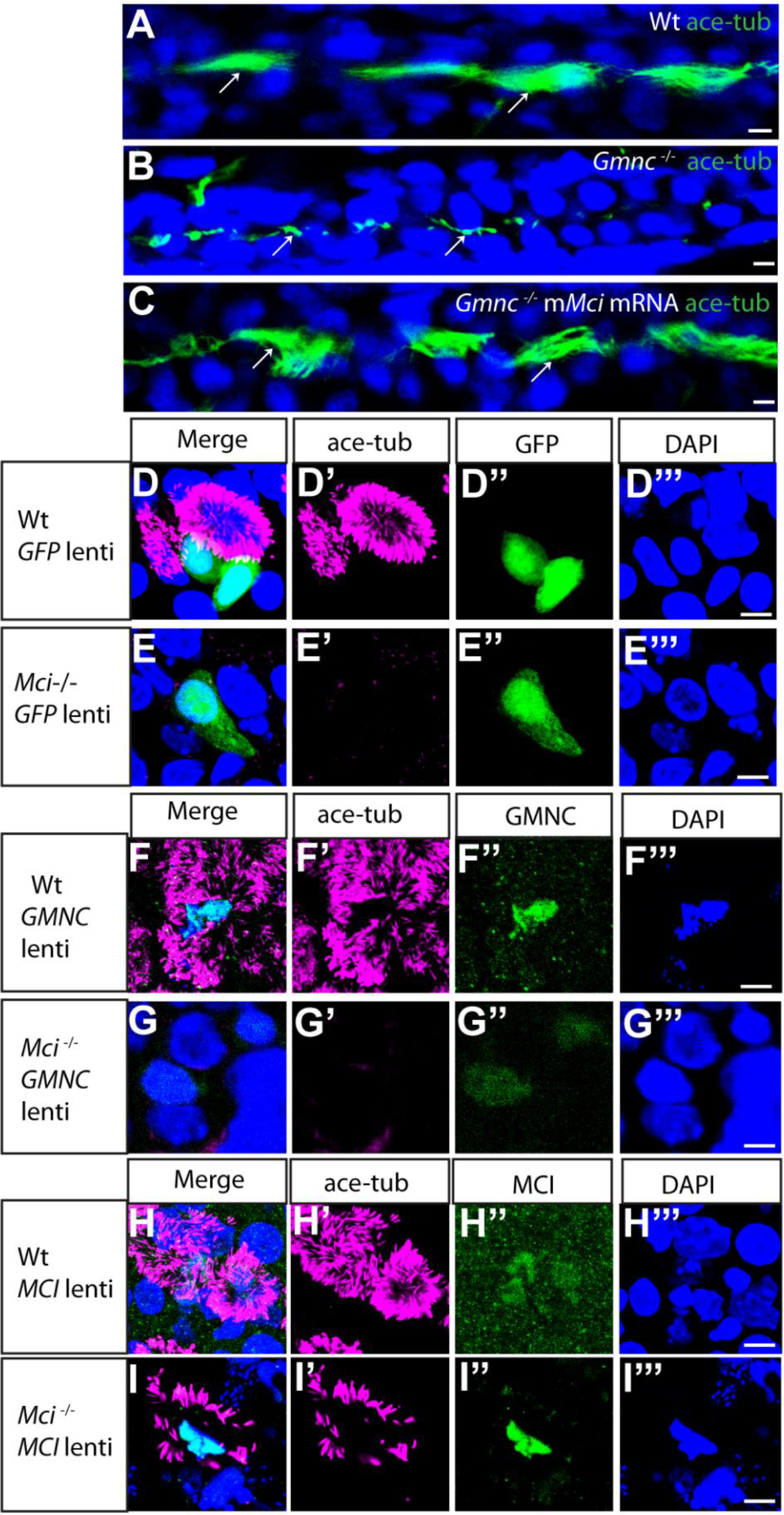
MCI can substitute for GMNC activity, but not *vice versa*, in MCC differentiation. (A) Pronephric duct of a 48 hours post-fertilization (hpf) wild-type zebrafish embryo showing multiple cilia on MCCs (arrows). (B) Pronephric duct of a 48 hpf *gmnc* mutant zebrafish embryo showing severe lack of MCCs. Monocilia, which are not affected by the loss of Gmnc, are indicated (arrows). (C) Pronephric duct of a 48 hpf *gmnc* mutant embryo showing rescue of MCCs (arrows) on over-expression of mouse *Mci* (m*Mci*) mRNA. 16 zebrafish embryos over-expressing mouse MCI were genotyped. 5 were *gmnc* homozygotes of which 3 showed MCC rescue in pronephric ducts (partial to full rescue in one or both ducts). (D) Lentivirus-mediated over-expression of GFP in wild-type airway cell ALI culture does not affect MCC differentiation (D’-D’’’ shows individual channels). (E) Over-expression of GFP in *Mci* mutant airway cell ALI culture does not restore MCC differentiation (E’-E’’’ shows individual channels). (F) Over-expression of GMNC in wild-type airway cell ALI culture induces supernumerary MCC differentiation (F’-F’’’ shows individual channels). (G) Over-expression of GMNC in *Mci* mutant airway cell ALI culture does not rescue MCC differentiation (G’-G’’’ shows individual channels). (H) Over-expression of MCI in wild-type airway cell ALI culture induces supernumerary MCC differentiation (H’-H’’’ shows individual channels). (I) Over-expression of MCI in *Mci* mutant airway cell ALI culture rescues MCC differentiation (I’-I’’’ shows individual channels). In all preparations, cilia were stained with anti-acetylated tubulin antibodies (green in a-c; magenta in d-i), and nuclei with DAPI (blue). Over-expressed GFP, GMNC and MCI were detected with anti-GFP and anti-HA antibodies, respectively (green in D-I). Lentivirus-mediated over-expression of GFP, MCI and GMNC in ALI cultures represents 2 independent biological replicates. Scale bars, 5 μm.

## DISCUSSION

Using *Mci* mutant mice, we have established two distinct steps in the developmental pathway for MCC formation that had remained previously unrecognized and are genetically separable: first, GMNC acts to specify MCC precursors, whereas in the second step, MCI drives multiple basal body production and multiciliation. Thus, in the absence of GMNC function, the MCC-specific developmental program is blocked at the earliest step, and no MCC precursors are generated (Zhou et al., 2015, Terré et al., 2016, Arbi et al., 2016). By contrast, loss of MCI does not derail MCC precursor specification, but affects their subsequent differentiation into MCCs. Although we cannot rule out species-specific differences in MCI function, it is likely that the discrepancy between our findings and the currently held notion of MCI activity (required for MCC specification and differentiation) stems from the different strategies used to interrogate MCI in mice and frogs (genetic mutant in mice versus morpholino knock-down in frogs) as well as methods used to examine MCC status on MCI loss in mice and humans (*in vivo* and *in vitro* analysis of MCCs from multiple mouse ciliated tissues versus RGMC patient MCCs obtained using nasal brush biopsy) (Stubbs et al., 2012, Boon et al., 2014).

Analysis of various kinds of multiciliated epithelia from *Mci* mutant mice have revealed that in all instances, MCC precursors form and express several transcription factors necessary for ciliary differentiation and motility. Consistent with this, these precursors differentiate into cells with a single motile-like cilium. However, we could not detect multiple basal bodies with several makers of these organelles as well as TEM analysis. Even though some deuterosomes do form, no mature basal bodies are ultimately generated. Indeed, expression of genes currently implicated in the DD pathway – *Deup1*, *Ccdc78*, *Ccno* and *Cdc20b* - is significantly reduced in the *Mci* mutants. Since some MCC basal bodies are thought to be produced via the MCD pathway (Al Jord et al., 2014), our observation that there is consistently only one basal body in *Mci* mutant MCCs suggests that this pathway is also strongly impaired. Yet, we did not detect major changes in the levels of several important MCD pathway genes. Lack of effect on the MCD pathway genes have also been reported previously for the *Gmnc* mutant mice (Terré et al., 2016). Moreover, since MCC precursors devoid of both mother and daughter centrioles can generate deuterosomes and multiple basal bodies (Zhao et al., 2018), these data and our findings can be taken to indicate that the MCD pathway may not function in MCCs at all, at least in the tracheal MCCs, which we have investigated in sufficient detail, and all of the basal bodies in these cells could arise exclusively via the DD pathway. Given all of the current ambiguity by which the deuterosomes and basal bodies arise in the MCCs (Al Jord et al., 2014, Zhao et al., 2018), the *Mci* mutant mice will be a valuable reagent for further investigations into the precise mechanisms involved in these processes.

Finally, we have provided biochemical evidence for the difference in the transcriptional activities of GMNC and MCI, which explains the distinct MCC phenotypes observed when they are individually mutated. We found that GMNC interacts much more efficiently with E2F5 than E2F4. In addition, replacement of the C-terminal portion of GMNC with that from MCI, conferred on the chimeric protein the ability to interact with E2F4. Furthermore, our data show that GMNC is more effective in inducing *FOXJ1* and *MCI*, whereas MCI is more effective in inducing genes involved in basal body generation. When they are over-expressed with E2F4 or E2F5, GMNC is able to induce its targets much more efficiently with E2F5, whereas MCI largely fares equally well with E2F4 and E2F5. These data suggest that GMNC, in association with E2F5, induces expression of *Mci* and *Foxj1* to generate MCC precursors, but does not activate genes for basal body production. MCI, being more promiscuous in its ability to interact with the E2F factors, then amplifies the expression of *Foxj1* (and genes for other ciliary transcription factors) for the massive upregulation of the motile cilia transcriptional program, but more importantly, induces genes for the production of multiple basal bodies. This molecular logic helps to clarify why GMNC cannot rescue MCC formation in *Mci* mutant ALI culture, but MCI is sufficient to restore MCC development in *gmnc* mutant zebrafish. Although the C-terminus is essential for conferring the differential interaction with E2F proteins, the N-terminal portion of GMNC appears to be equally important for its transcriptional ability. The chimeric GM protein not only failed to mimic the transcriptional activation profile of MCI, but also showed an overall impairment in transcriptional activating activity. This implies that either the N-terminal is important for interacting with other transcriptional cofactors (since the coiled coil domain resides in this region) or it makes an important contribution to the formation of a functional E2F/DP1 ternary complex. As a corollary of this observation, we propose that the N-terminus of MCI could also have a similar role in determining its transcriptional activity. Since the precise mechanism by which the MCI/GMNC-E2F-DP1 complex regulates transcription is not understood, and it is also not clear whether other co-factors are involved (especially in the regulation of the distinct sets of target genes), further biochemical experiments will be required to resolve these questions.

In conclusion, our study of the *Mci* mutant mouse will be of direct relevance to the role of MCCs in ciliopathies, especially for the pathobiology of RGMC, a relatively new but acute airway disease that remains rather poorly defined. In addition, the ability of GMNC and MCI to generate ectopic MCCs provides a powerful avenue to devise strategies for restoration of functional ciliated epithelia by gene therapy. This holds promise not only for rare disorders like RGMC, but also in acquired and more prevalent airway pathologies such as chronic obstructive pulmonary disorder (COPD), where impairment of ciliary function has also been implicated (Yaghi and Dolovich, 2016).

## MATERIALS AND METHODS

### Ethics approvals

All mouse and zebrafish experimentation was performed under approval from the Singapore National Advisory Committee on Laboratory Animal Research and conformed to the stipulated ethical guidelines.

### Generation of *Mci* knockout mice

*Mci* mutant mice were generated by CRISPR/Cas9 mediated deletion of a DNA fragment within exon 2 of the *Mci* gene. Two guide RNAs (gRNAs) were designed to target the exon 2, and were co-injected with Cas9 mRNA (25 ng/μl) into C57BL/6 one-cell embryos at a concentration of 15 ng/μl each (see Table EV1 for sequences of gRNAs and all primers used in this study). A total of 247 embryos were injected, out of which 130 were implanted into 9 pseudo-pregnant females. Founder animals were screened by PCR, and mutations were determined first by T7 endonuclease I assay, and then by deep sequencing of PCR products (for selected founders). Out of 13 pups born alive, 9 were found to contain mutations at the *Mci* targeted region. Founders containing the desired mutation were bred with the wild type C57BL/6J animals to produce F1 heterozygotes. The F1 mutants were identified by PCR and confirmed by sequencing.

### Zebrafish strains

The AB strain was used as the wild-type for all experiments. The *gmnc* mutant strain has been described previously (Zhou et al., 2015).

### DNA constructs

Coding sequences for human and mouse DP1, E2F4, E2F5 were cloned into the pCS2 vector with 6x Myc-tags at the N-terminus. Coding sequences for human and mouse GMNC and MCI were cloned into the pXJ40 vector with one HA tag at the N-terminus. The human GM chimera was generated using overlapping extension PCR, and cloned into pXJ40 vector with one HA tag at the N-terminus.

### Co-immunoprecipitation and immuno-blot

Desired combinations of plasmids were co-transfected into HEK293T cells, in 10 cm dishes (3 μg per plasmid, per dish) using Lipofectamine 2000 (Thermo Fisher Scientific). After 24 hrs of incubation, transfected cells were lysed in 800 μl of RIPA buffer (Thermo Fisher Scientific) supplemented with complete Mini protease inhibitors, EDTA-free (Roche, #11836170001). The cell lysates were sonicated briefly and spun down. An aliquot was taken from the clear cell lysate and boiled in 1X SDS loading buffer as input (TCL), and the rest was rotated over-night with 25 μl of Protein A-agarose beads (Roche) and 2 μg of mouse anti-HA antibody (monoclonal, Santa Cruz, SC7392). The beads were washed four times in the IP buffer and boiled in 50 μl of 1X SDS loading buffer (IP:HA). Both TCL (15 μl, 1 %) and IP (15 μl, 30 %) were resolved by SDS-PAGE gels, transferred to PVDF membranes, blocked in 2 % BSA, and probed with relevant primary antibodies (rabbit anti-HA (Santa Cruz, SC805); rabbit-anti-Myc (Santa Cruz, SC289) and secondary antibodies (anti-mouse HRP conjugate (Promega, #W4028), anti-rabbit HRP conjugate (Promega, #W4018)).

### Antibodies

Primary antibodies: mouse-anti-HA (Santa Cruz SC7392, 1:2500 for western blot, 1:500 for immunofluorescence (IF)); rabbit-anti-HA (Santa Cruz SC805, 1:2500 for western blot, 1:500 for IF); mouse-anti-Myc (Santa Cruz SC40, 1:2500 for western blot); rabbit-anti-Myc (Santa Cruz SC289, 1:2500 for western blot); mouse-anti-acetylated-α-tubulin (Sigma-Aldrich T 6793, 1:500 for IF); mouse-anti-α-tubulin (Sigma-Aldrich T6557, 1:500 for IF); mouse-anti-γ-tubulin (Sigma-Aldrich T6557, 1:500 for IF); rabbit-anti-γ-tubulin (Sigma-Aldrich T5192, 1:500 for IF); mouse-anti-γ-tubulin (Sigma-Aldrich T6557, 1:500 for IF); rabbit-anti-RFX2 (Sigma-Aldrich HPA048969, 1:250 for IF), rabbit-anti-RFX3 (Sigma-Aldrich HPA035689, 1:250 for IF); rabbit-anti-FOXJ1 (Sigma-Aldrich, HPA 005714, 1:250 for IF); mouse anti-FOXJ1 (ebiosciences 14-9965-80, 1:100 for IF); rabbit-anti-TAP73 (Abcam ab40658, 1:250 for IF); rabbit-anti-RSPH1 (Sigma-Aldrich HPA017382, 1:250 for IF); rabbit-anti-RSPH9 (Sigma-Aldrich HPA031703, 1:250 for IF); rabbit-anti-PERICENTRIN (Abcam ab4448, 1:250 for IF); mouse anti-CENTRIN (EMD Millipore Corp 04-1624, 1:200 for IF) and rabbit-anti-DEUP1 (kind gift of X. Zhu, Shanghai Institute of Biochemistry and Cell Biology, 1:200 for IF). Secondary antibodies (all used at 1:500 for IF): Alexa 488 goat-anti mouse (Invitrogen A-11029); Alexa 488 goat anti-rabbit (Invitrogen A-11034); Alexa 555 goat anti-rabbit (Invitrogen A-21428); Alexa 555 goat anti-mouse (Invitrogen A-28180).

### Cell and ALI culture

HEK293T and HEK293FT cells were cultured in DMEM with 4500 mg/l glucose and 10 % FBS (HyClone, SH30071.03HI). mTEC culture was performed according to published protocol (Vladar and Brody, 2013). Briefly mTEC cells were grown on transwells with transparent PET membrane (Life Science, 353095) in mTEC plus+RA medium (DMEM/F12; Life Science, 11330-032), Fungizone (Life Technologies, 15290-018, 0.1 % v/v), Insulin (Sigma-Aldrich, l1882, 10 mg/ml), Epidermal growth factor (BD Biosciences, 354001, 25 ng/ml), Transferrin (Sigma T1147, 5 mg/ml), Cholera toxin (Sigma-Aldrich C8052, 0.1 mg/ml), Fetal bovine Serum (Life Technologies 26140-079, 5% v/v), ROCK inhibitor (ATCCY27632, 10 μM), Retinoic acid (Sigma-Aldrich R2625, 50 nM) and Penicillin-Streptomycin (Life Technologies 15140-148, 100 U Pen, 100 mg Strep per ml). When cells on the apical side of the transwell chambers reached 100 % confluence, ALI was established by aspirating the culture medium from the transwell chambers, and addition of differentiation medium (mTEC Plus medium without fetal bovine serum and ROCK inhibitor) to the basal chamber on 24 well-plates. The mTEC cells were maintained on transwells by changing the differentiation medium in the basal chamber every 2 days.

### Immunofluorescence

For IF analysis, mTEC cells grown on transwells were fixed in 4 % paraformaldehyde (PFA) at room temperature (RT) for 10 minutes and permeabilised with PBTX (PBS, 0.5 % Triton X-100) for 2 hrs and washed in phosphate buffered saline (PBS). Cells were then blocked with 2 % bovine serum albumin in PBS for 1 hr, followed by 1 hr with primary antibody at RT. After 3 washes in PBS, cells were incubated with secondary antibodies and DAPI for 1 hr. After briefly washing with PBS, the cells were mounted on glass slides with fluorescence mounting medium and imaged using an Olympus FluoView upright laser scanning confocal microscope. Cryosections of mouse tissues were prepared by the histopathology unit and the slides stored at −80°C. On the day of staining, slides were thawed and dried before drawing borders around the sections with a PAP pen (Abcam ab2601). The slides were then fixed with 4% PFA for 15 min at RT in Coplin jars (all subsequent steps performed in Coplin jars unless stated otherwise). They were rinsed twice with cold PBS followed by permeabilisation with 0.2 % Triton (in PBS) for 15min. They were washed 3 times, 5 min each in PBS and blocked with 2 % BSA in PBS for 2 hrs at RT. The slides were then transferred to a humidified box. Primary antibodies in PBS (with 0.1 % Tween20 and 1 % BSA) were pipetted onto the sections and incubated over-night at 4°C. The following day, slides were washed with PBS on a shaker, 6 times, 10 min each, at RT. Secondary antibodies in PBS (with 0.1 % Tween20 and 1 % BSA) were then added, and the slides incubated in the humidified box for 5 hrs at RT. Finally, the slides were washed 6 times (10 min each) with PBS at RT, dried, and mount with Vectashield.

### RT-qPCR

cDNA preparations were generated using the SuperScript III First-Strand Synthesis System (Invitrogen 18080051). Gene-specific primers for qPCR were designed using the Primer3 software (Primer3 (v.0.4.0)) and are listed in Table EV1. qPCRs were performed with the EXPRESS SYBR GreenER Super Mix (Invitrogen A10315) on an Applied BioSystems 7900HT Fast Real-Time PCR System using the SDS2.4 software. Technical triplicate reactions were performed for each sample. Using Microsoft Excel, gene expression fold differences were calculated from the Ct values after normalizing against the internal control *Gapdh*/*GAPDH*.

### Microinjection of zebrafish eggs and processing for IF analysis

mRNA encoding mouse MCI (300 ng/μl, 0.75 nl per egg) was injected into one cell stage eggs derived from in-cross of *gmnc* heterozygous fishes. At 48 hours post fertilization (hpf), the injected embryos were fixed with Dent’s fixative (80 % methanol, 20 % DMSO) for 3 hrs at RT and then subjected to IF staining using routine protocol.

### Lentivirus generation and infection

Gene expression lentiviruses were generated using ViraPower™ Lentiviral Expression Systems Version C (Invitrogen 25-0501). Briefly, coding sequences of different genes were cloned into PLVX vector, followed by transfection into HEK293FT cells together with the Lentiviral Packaging Mix (Invitrogen, K4975-00). Viruses were harvested by collecting the cell culture medium 3 days after transfection. Viral titration was performed by infecting 293FT cells with the control GFP lentivirus which was generated together with gene-specific expression lentiviruses (*GMNC* and *MCI*), and then determined by the percentage of GFP positive cells 3 days after infection. For confluent mTEC cells viral infection, the cells were treated with 12 mM EGTA (Sigma-Aldrich, E3889) in 10 mM HEPES (Sigma-Aldrich H3375-25G), pH 7.4 for 25 min at 37°C. After washing the EGTA treated cells with PBS, a mix of specific amounts of lentivirus and Polybrene (Sigma-Aldrich, H9268, 5 μg/ml final concentration) was added into the culture medium. mTEC cells with the viruses were then centrifuged at 1,500g for 80 min at 32oC, and grown at 37oC in a cell culture incubator.

### Electron microscopy of mouse trachea

For SEM analysis: Immediately after dissection, mouse tracheae were fixed by immersion in 4 % formaldehyde and 2 % glutaraldehyde (EM grade, Electron Microscopy Sciences) in 0.1M Sodium cacodylate buffer (pH = 7.4) for 12 hrs. After washing, samples were cut across the length into approximately 2 mm pieces, and subsequently cut longitudinally to expose the interior surface. Trimmed samples were post-fixed with 1 % Osmium tetroxide in distilled water for 2 hrs, washed with distilled water and dehydrated in Ethanol series. Dehydrated samples were dried using critical point drying (Leica EM CPD030), mounted onto aluminium stubs with trachea lumen facing up and sputter coated with 4 nm layer of platinum (Leica EM SCD050). SEM analysis was performed using a JSM 6701F SEM (JEOL) microscope operating at 2.5kV. Images were collected from random areas of the wild-type and mutant samples. For TEM analysis: Dissected tracheae were fixed in 4 % paraformaldehyde, 2.5 % glutaraldehyde, and 0.2 % picric acid in 0.1 M Sodium cacodylate buffer. Samples were washed in Sodium cacodylate buffer and post fixed with 1 % Osmium tetroxide.

Samples were again washed in Sodium cacodylate buffer before dehydration through a graded series of Ethanol. After dehydration, samples were infiltrated and embedded with Spurr resin (Electron Microscopy Sciences 14300) before polymerization at 60°C. Ultra-thin sections were obtained by cutting sample blocks on an ultramicrotome (Leica ultracut UCT), stained with 4 % Uranyl acetate and 2 % Lead citrate before viewing the sections with a TEM (Jeol 1010) microscope.

### Statistical analysis

The statistical analysis, including standard error of the mean (SEM), standard deviation (SD), and unpaired *t*-test, was performed using the software GraphPad Prism 7.04.

## Acknowledgements

We thank the Animal Gene Editing Laboratory, Biological Resource Centre, Agency for Science, Technology and Research for generating the *Mci* mutant mice, the Advanced Molecular Pathology Laboratory for histological services, the Institute of Medical Biology-Institute of Molecular and Cell Biology Joint Electron Microscopy Suite for electron microscopy analysis, V. Tergaonkar for assistance in obtaining appropriate clearance from the Institutional Animal Care and Use Committee (IACUC) for experiments with the *Mci* mutant mice, X. Zhu for DEUP1 antibodies and A. Guha and members of our laboratory for discussion and comments on the manuscript.

## Author contributions

S.R. conceived the project. L.H. performed majority of the experiments including analysis of mutants, ALI culture and transcriptional studies. P.A. established ALI culture and gene expression analysis. F.Z. contributed to mutant analysis. Y.L.Z. contributed data for protein interaction studies. Y.L.C. performed TEM analysis. S.R. and C.D.B. supervised the work. All authors critically analyzed the data. L.H. and Y.L.Z. assembled the figures. S.R. wrote the paper with input from all authors.

## SUPPLEMENTAL INFORMATION

**Fig. S1.**
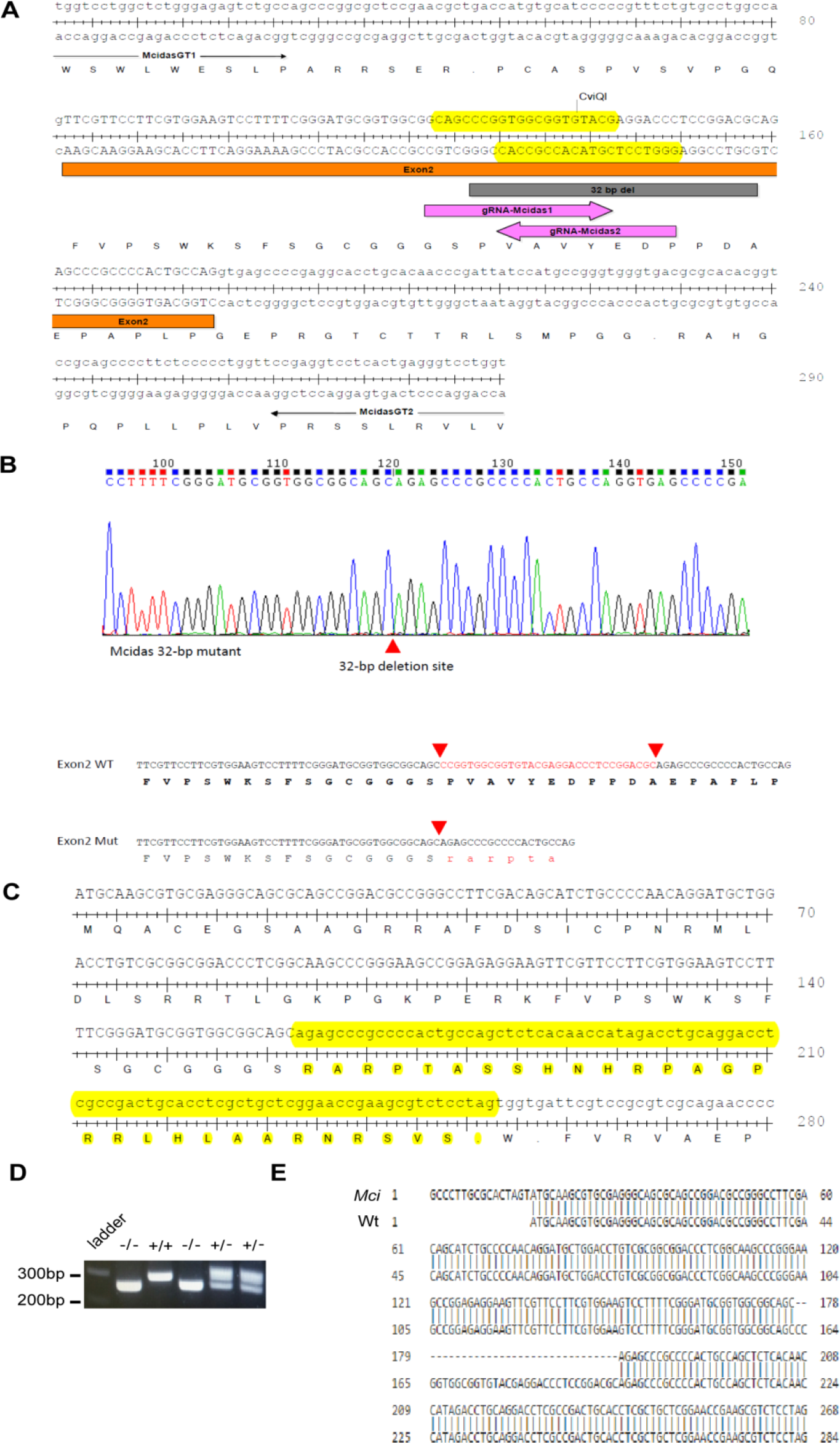
Generation of a deletion allele at the mouse *Mci* locus. (A) Partial genomic sequence of the mouse *Mci* gene, showing the gRNAs (pink arrows) and their target sites on the forward and reverse strands (highlighted in yellow) used to induce a 32 bp deletion within exon 2. Binding sites for genotyping primers (McidasGT1 and McidasGT2) are also indicated. (B) Electropherogram showing 32 bp deletion in *Mci* exon 2. Also shown below is the conceptual translation of the wild-type and mutant *Mci* coding sequence around the deletion site. (C) Conceptual translation of the predicted mutant *Mci* ORF shows a highly truncated MCI protein, retaining only 54 native amino acids at the N-terminus. Sequences highlighted in yellow indicate disruption of the reading frame before the premature STOP codon. (D) Gel image of DNA fragments amplified in wild-type, heterozygote and homozygous *Mci* mutants using primers flanking the 32 bp deletion. Size of the wild-type band is 290 bp and the mutant band is 258 bp. (E) Sequence analysis of *Mci* cDNA obtained from tracheal tissue of the homozygous mutants confirms a deletion of 32 bp.

**Fig. S2.**
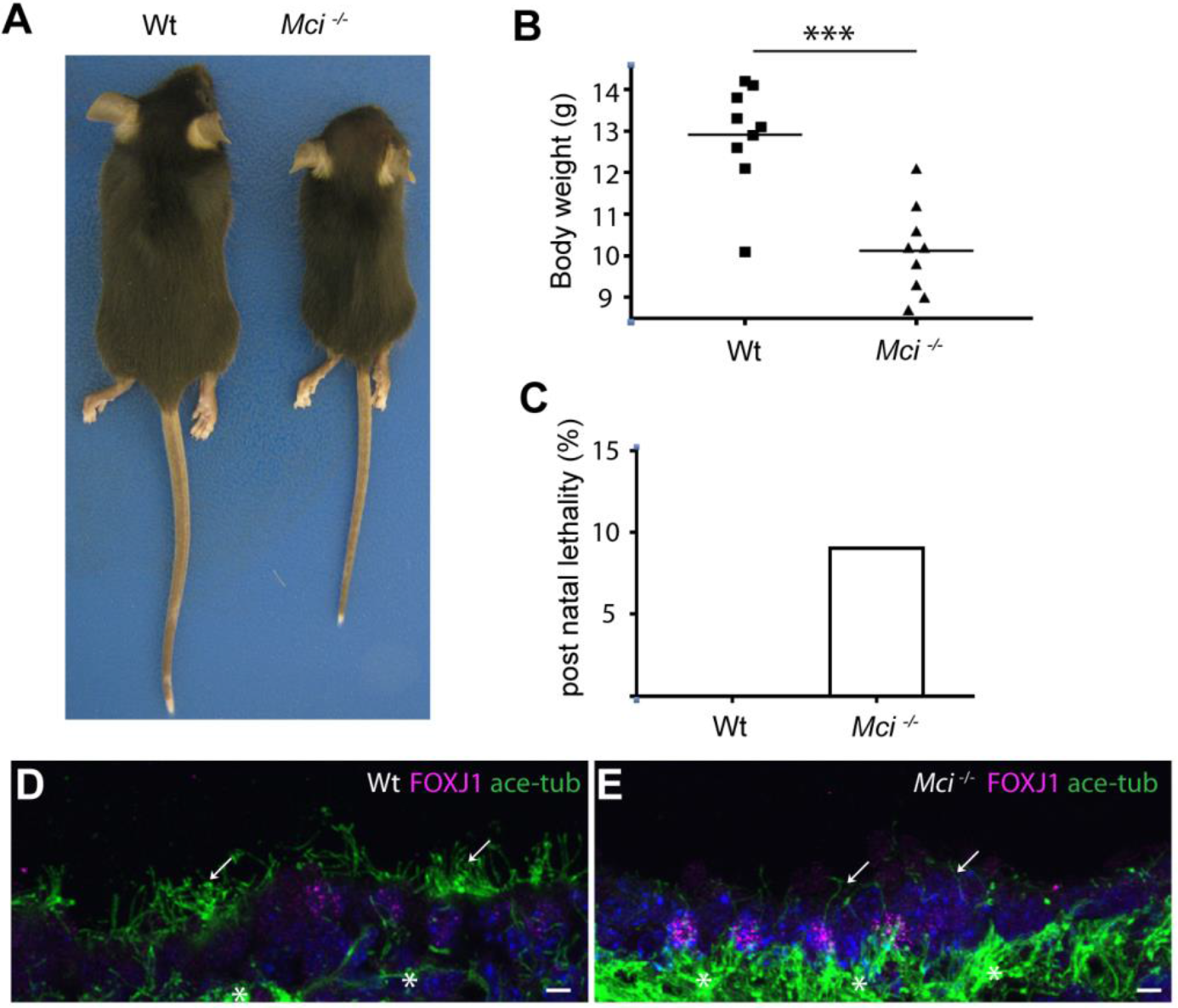
Gross phenotypes of *Mci* knockout mice. (A*) Mci* knockout mice are smaller in size compared to the wild-type. (B) The body weight comparison between wild-type and *Mci* mutant mice at post-natal day (P) 28. *n* = 9 for each genotype. (C) Percentage of lethality of wild type and *Mci* knockout mice at P28. *n* = 22 for each genotype. (D) Nuclear localized FOXJ1 expression in MCCs of wild-type brain ependyma. Multicilia are indicated by arrows and the cytoskeletal microtubule network by asterisks. (F) Nuclear localized FOXJ1 expression in monociliated cells of *Mci* mutant brain ependyma. Monocilia are indicated by arrows and the cytoskeletal microtubule network by asterisks. Scale bars, 5 μm.

**Fig. S3.**
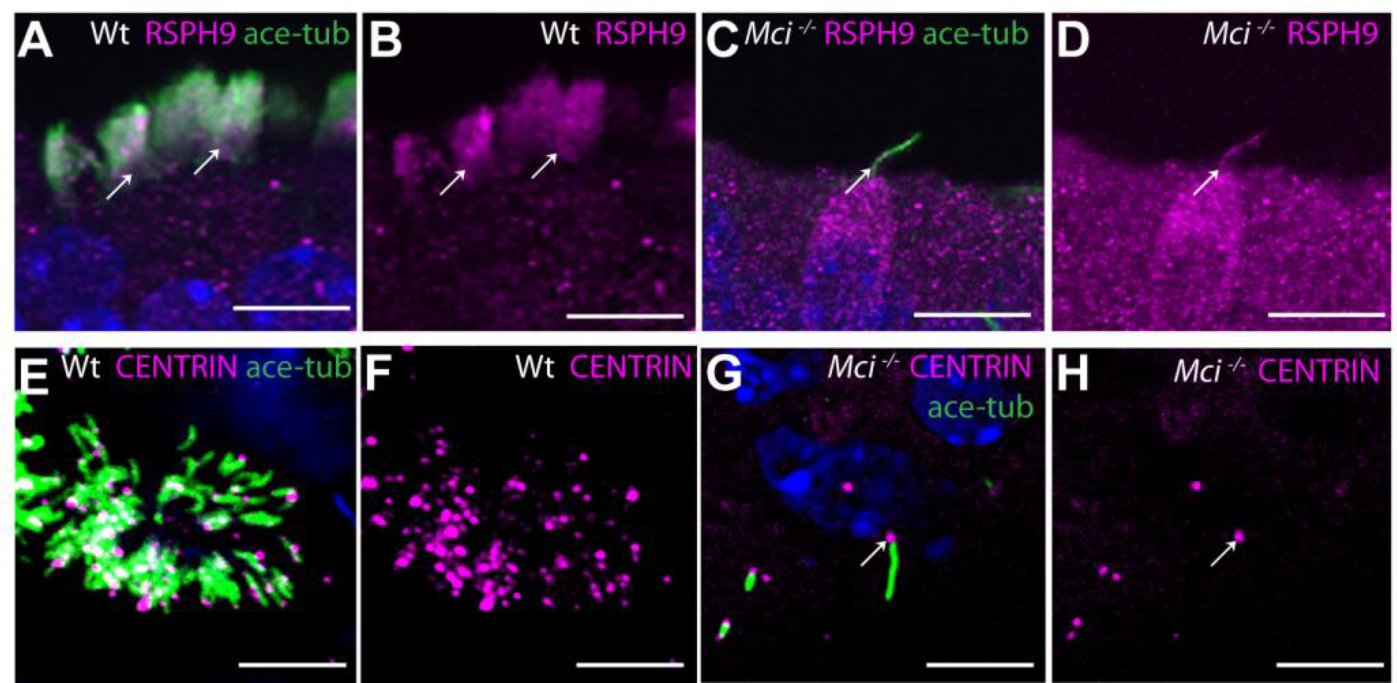
*Mci* mutant MCCs precursors differentiate a single cilium that localizes motile cilia-specific proteins but are unable to make multiple basal bodies. (A) RSPH9 co-localization with acetylated tubulin to MCC cilia of wild-type trachea (arrows). (B) RSPH9 localization to MCC cilia of wild-type trachea (arrows; display of only RSPH9 staining from panel A). (C) RSPH9 co-localization with acetylated tubulin to single cilium of *Mci* mutant trachea (arrow). (D) RSPH9 localization to single cilium of *Mci* mutant trachea (arrow; display of only RSPH9 staining from panel C). (E) Wild-type MCC differentiated in ALI culture with multiple basal bodies (stained with anti-CENTRIN antibodies) and multiple cilia. (F) Display of only CENTRIN staining from panel E. (G) *Mci* mutant cells differentiated in ALI culture with single basal body (expressing CENTRIN, arrow) and single cilium. (H) Display of only CENTRIN staining from panel G showing single basal body (arrow). In all preparations, cilia were stained with anti-acetylated tubulin antibodies (green) and nuclei with DAPI (blue). Scale bars A-D = 10 μm; E-H = 5 μm.

**Fig. S4.**
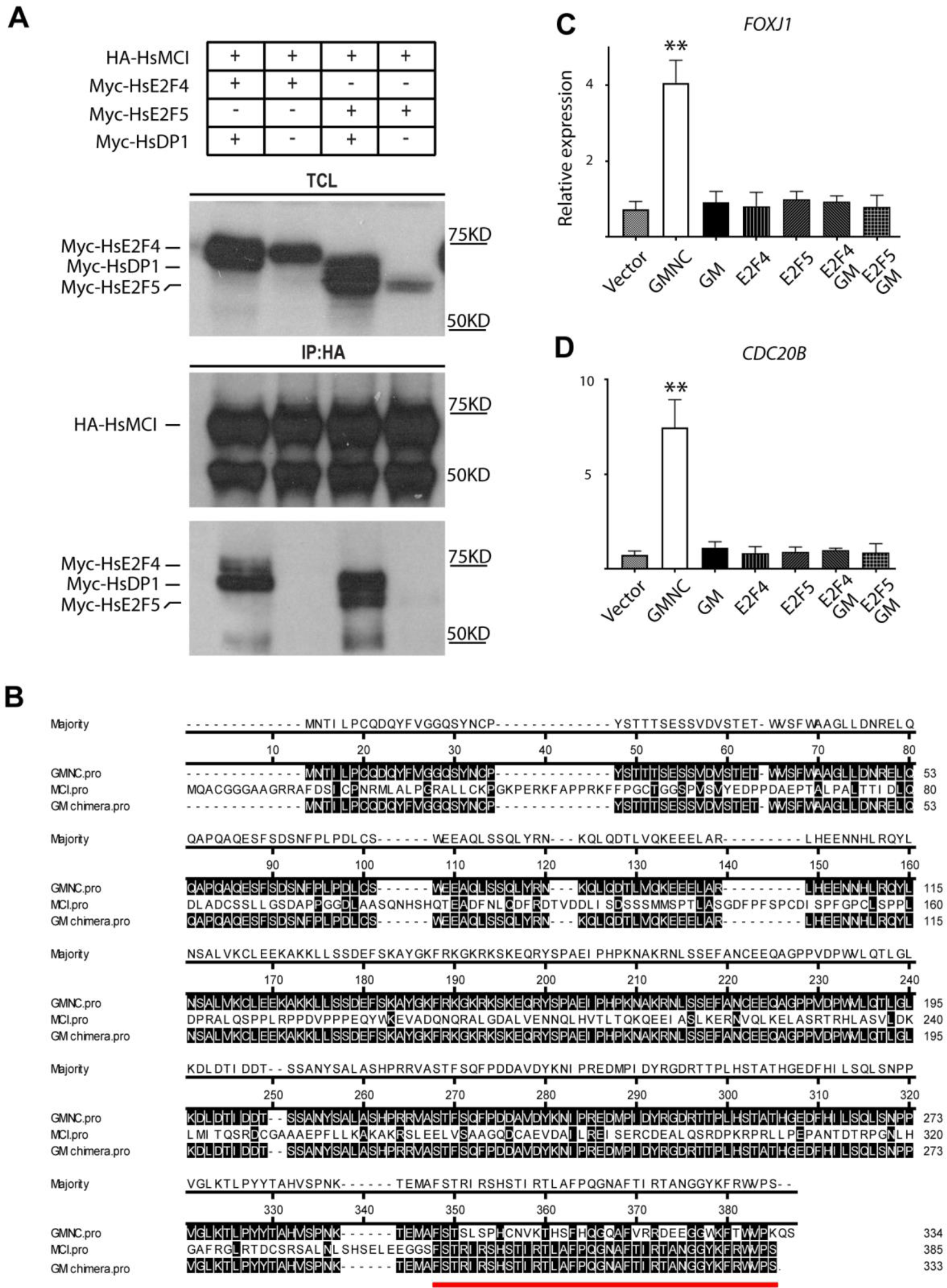
Interaction of MCI with E2F factors and transcriptional activity of the GMNC-MCI chimeric protein in HEK293T cells. (A) Co-immunoprecipitation data showing interaction of MCI with E2F4 as well as E2f5. Human proteins were used for this experiment. (B) Amino acid sequence alignment of human GMNC, MCI and GM proteins. The C-terminal fragment from MCI used to generate GM is underlined in red. (C) Unlike wild-type GMNC, the GM chimeric protein is unable to induce *FOXJ1* expression by itself or together with the E2F factors. (D) The GM protein is not more efficient in inducing *CDC20B* expression than wild-type GMNC either by itself or with the E2F factors. For C and D, relative expression levels have been plotted along the *y*-axis, and over-expression conditions indicated along the *x*-axis. Error bars: SEM. Immunoblot and qPCR data are representative of 2 independent biological replicates. p: ** ≤ 0.01.

**Fig. S5.**
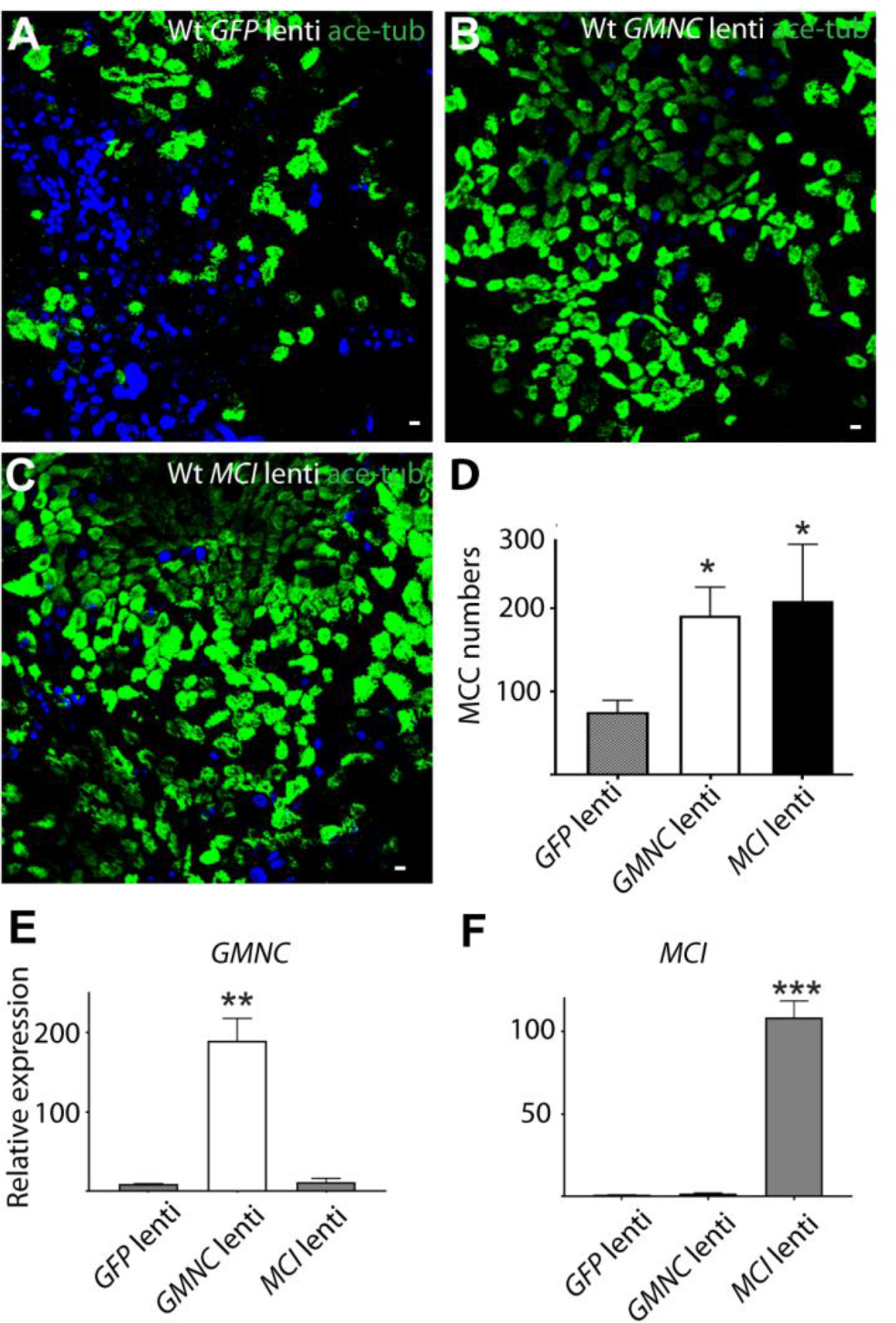
Over-expression of GMNC and MCI in wild-type airway cell ALI culture induces supernumerary MCCs. (A) Lentivirus mediated over-expression of GFP in wild-type airway cell ALI culture does not affect numbers of differentiating MCCs. (B) Over-expression of GMNC in wild-type airway cell ALI culture induces supernumerary MCCs. (C) Over-expression of MCI in wild-type airway cell ALI culture induces supernumerary MCCs. Scale bars, 5 μm. (D) Quantification of MCC numbers per field of view upon over-expression of GFP, GMNC and MCI in wild-type airway cell ALI cultures. (E,F) RT-qPCR analysis of GMNC and MCI expression levels on over-expression of GMNC and MCI in *Mci* mutant airway cells cultured under ALI conditions. Relative expression levels have been plotted along the *y*-axis, and over-expression conditions indicated along the *x*-axis. Lentivirus-mediated over-expression of GFP, MCI and GMNC in ALI cultures represent 2 independent biological replicates; qPCR analysis represents 2 independent technical replicates. Error bars: SEM. p: * ≤ 0.05, ** ≤ 0.01, *** ≤0.001.

**Fig. S6.**
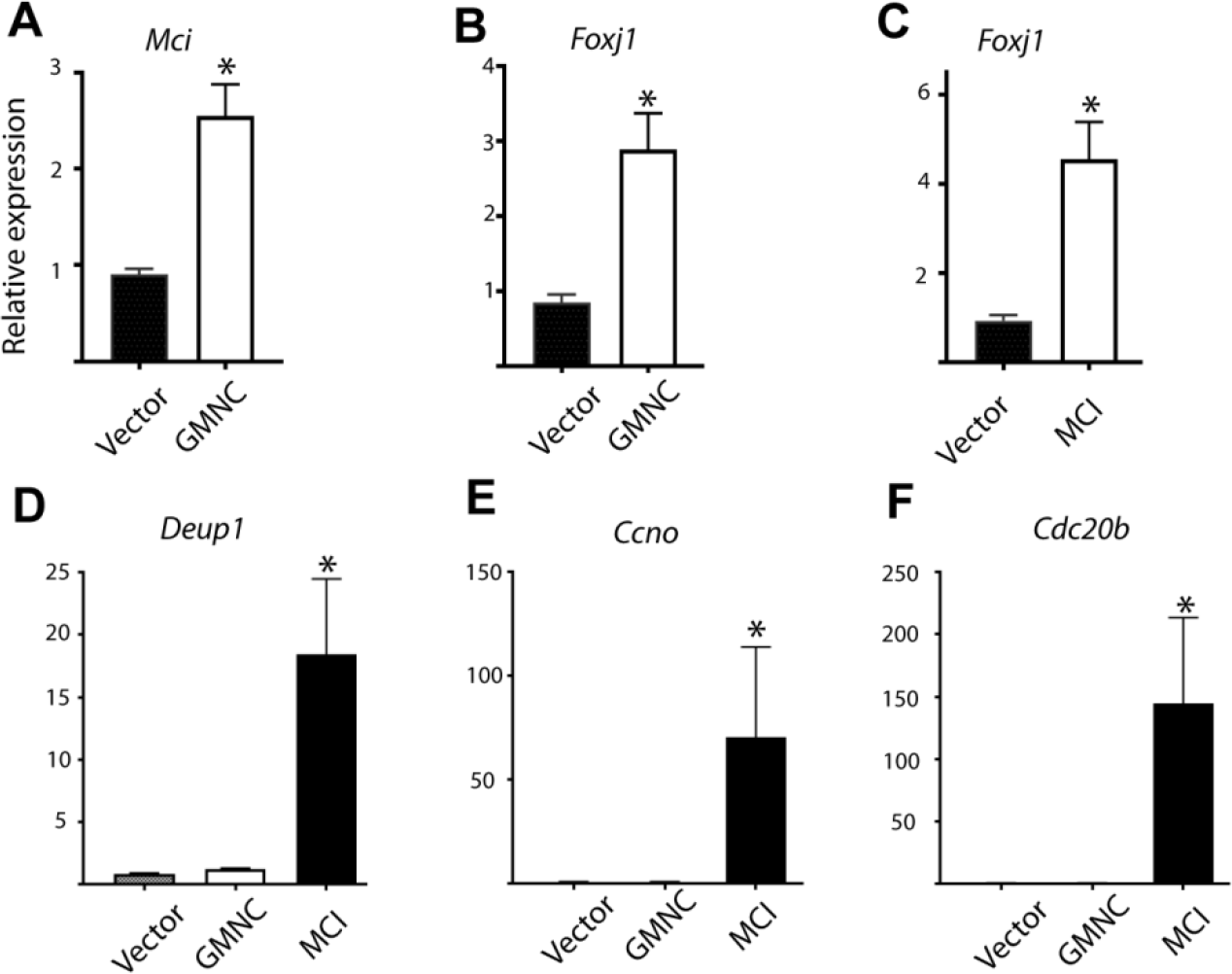
RT-qPCR analysis of ciliary transcription factor and DD pathway genes expression levels on over-expression of MCI and GMNC in *Mci* mutant airway cells cultured under ALI conditions. (A-F) Relative expression levels have been plotted along the *y*-axis, and over-expression conditions indicated along the *x*-axis. Error bars represent SEM. Analysis was done on 3 independent biological replicates. p: * ≤ 0.05.

**Table S1.**
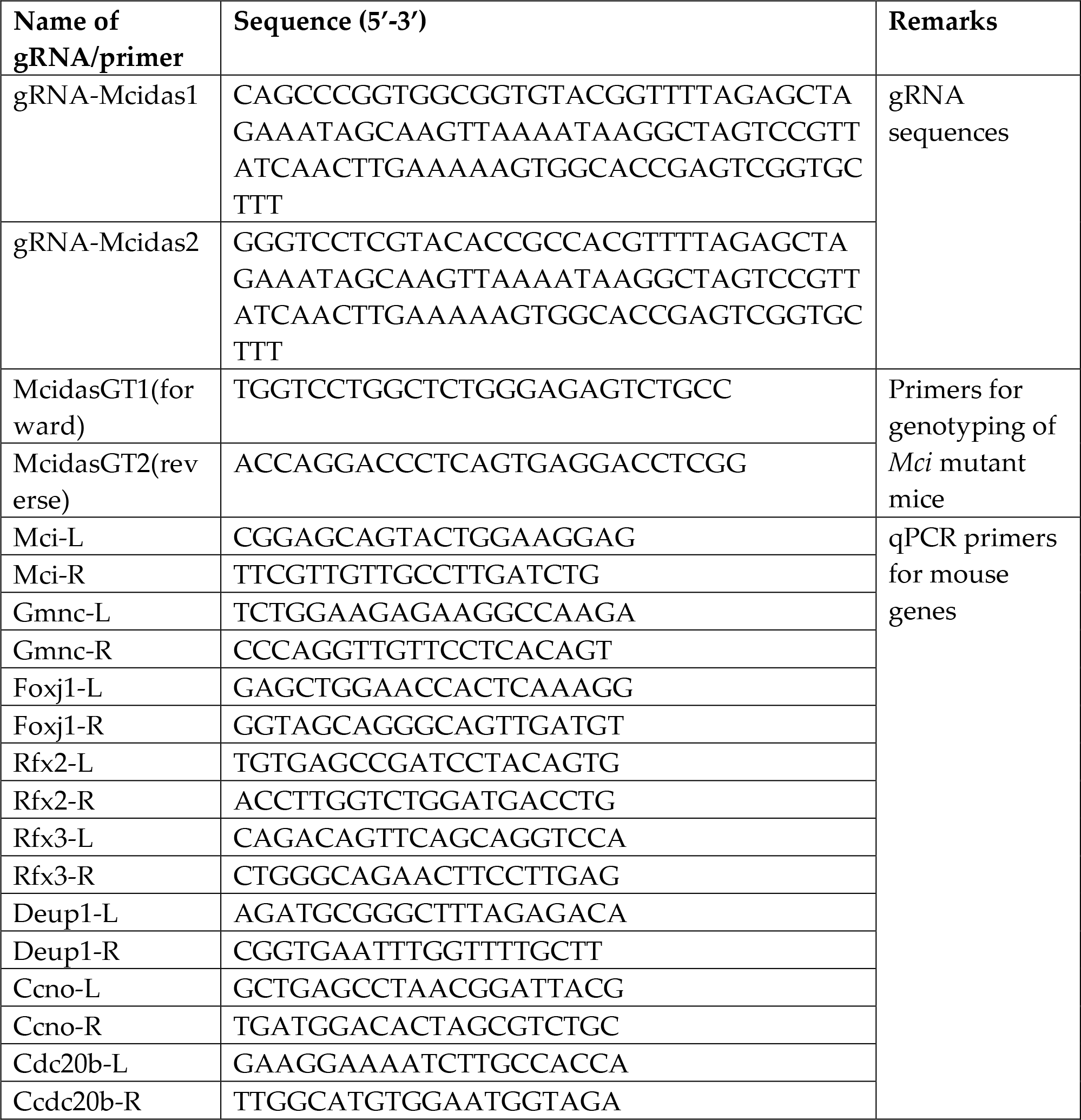
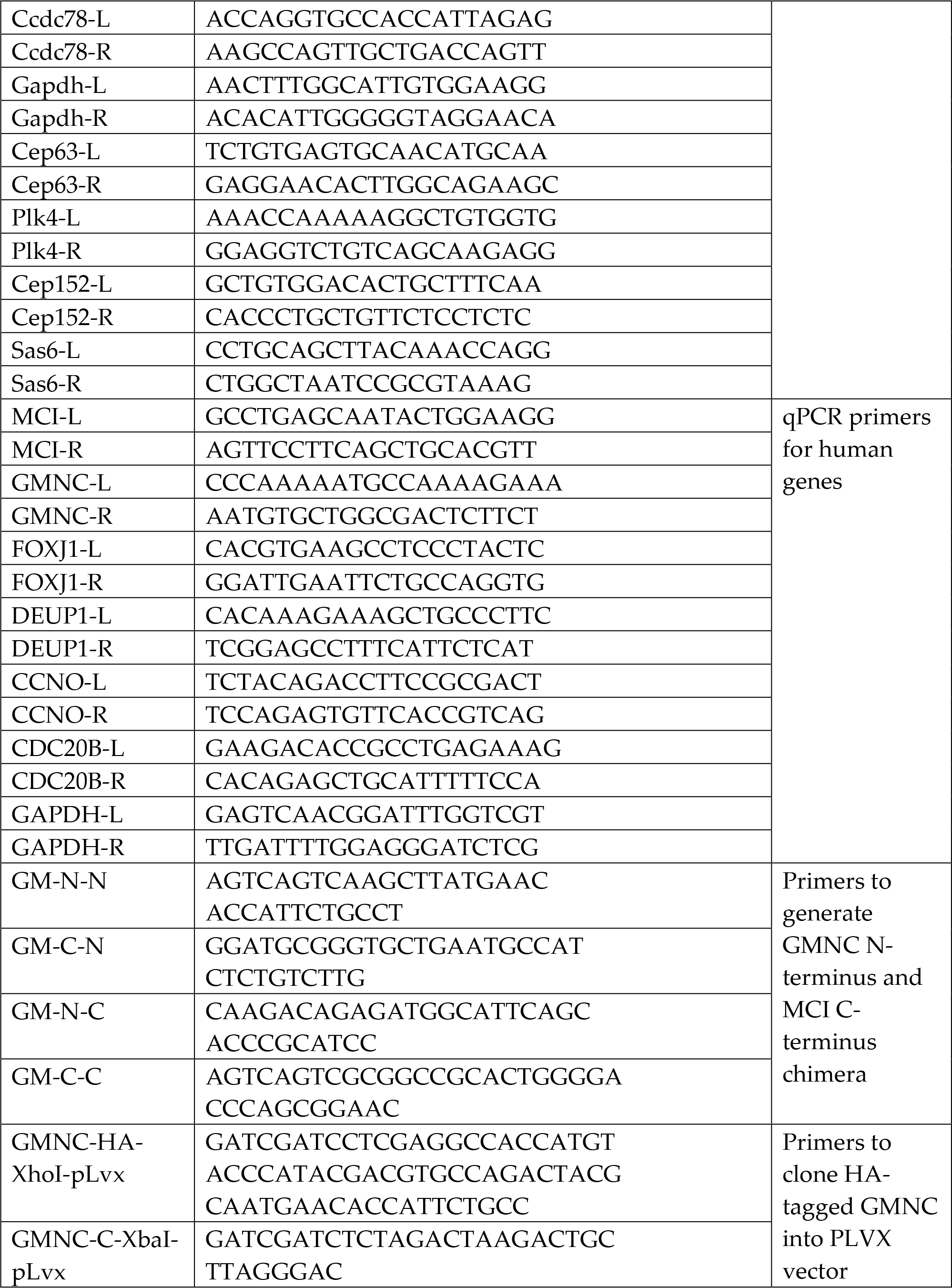
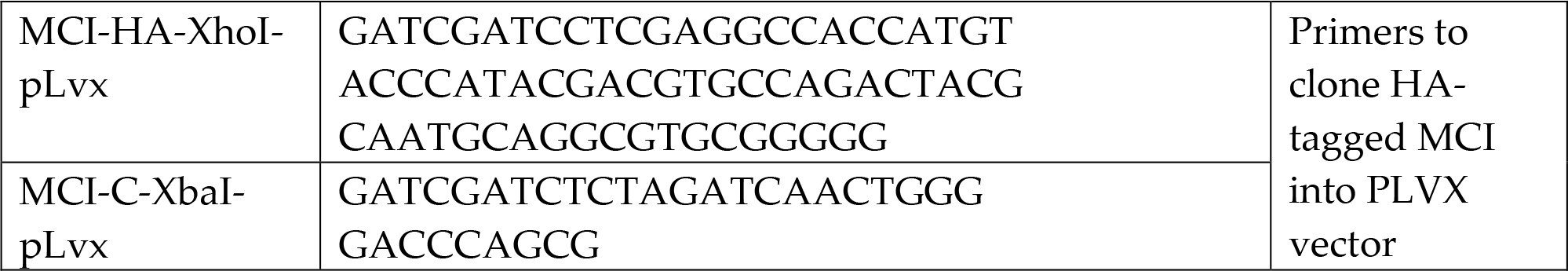

